# Bacterial aggregation triggered by low-level antibiotic-mediated lysis

**DOI:** 10.1101/2023.11.11.566689

**Authors:** Sharareh Tavaddod, Angela Dawson, Rosalind J Allen

## Abstract

Suspended bacterial aggregates play a central role in ocean biogeochemistry, industrial processes and probably many clinical infections – yet the factors that trigger aggregation remain poorly understood, as does the relationship between suspended aggregates and surface-attached biofilms. Here we show that very low doses of cell-wall targeting antibiotic, far below the minimal inhibitory concentration, can trigger aggregation of *Escherichia coli* cells. This occurs when a few cells lyse, releasing extracellular DNA – thus, cell-to-cell variability in antibiotic response leads to population-level aggregation. Although lysis-triggered aggregation echoes known trigger mechanisms for surface-attached biofilms, these aggregates appear to be of a distinct type, since they do not show typical biofilm characteristics such as protection from antibiotics, or increased biofilm-forming potential. Our work contributes to understanding the nature of bacterial aggregates and the factors that trigger their formation, and the consequences of widespread low-dose antibiotic exposure in the environment and in the body.

## Introduction

Many microbes form aggregates in liquid suspension. Aggregation can alter microbial physiology, enhancing antibiotic tolerance and biofilm formation^1–5^. It can also have important clinical, biophysical and ecological effects ^2,6,7^, including aggregate sinking that drives sequestration of carbon in the ocean and in wastewater treatment plants, preferential expulsion from the gut^8^ and loss of ability to invade a host^9^. Microbial aggregates have even been proposed as an early form of multicellular life^10^. However, despite its ubiquity and importance, in most cases the environmental factors that trigger aggregation, and the mechanisms by which they act, remain unclear.

In this work, we use *in vitro* experiments with *Escherichia coli* to show that bacterial aggregation can be triggered by exposure to antibiotic at very low doses (many times lower than the minimal inhibitory concentration). Our work supports previous reports linking low-dose antibiotic treatment with aggregation in aquatic bacterial communities^11^ and for *in vivo* gut bacteria in a zebrafish model^8^. The finding is significant because microbial exposure to low-dose antibiotics is widespread – occurring in wastewater, rivers and lakes, as well as during antibiotic therapy, agricultural use of antibiotics or due to ecological interactions between microbes (e.g. in soil). Low-dose antibiotic exposure is already a cause of concern because of its potential to enrich for antibiotic-resistant mutants^12^ or to trigger quorum sensing, virulence and biofilm formation^12–15^. Our work strengthens the case that it can also trigger aggregation, with associated biophysical and ecological consequences.

We also investigate the mechanism behind antibiotic-triggered aggregation of liquid-phase cells. It is well-known that microbial aggregation can be mediated by diverse mechanisms, including exopolysaccharides, extracellular DNA (eDNA) or proteins, and chemotaxis^1,4,16–21^. For antibiotic-triggered aggregation, our work reveals a central role for eDNA as the cohesive agent. We link the release of cohesive eDNA to the antibiotic-mediated lysis of a small subpopulation of bacteria. Since the vast majority of bacteria do not lyse at these very low antibiotic concentrations, it appears that the stochastic lysis of a few atypically antibiotic-sensitive cells within the population can drastically alter the fate of the population as a whole.

Our work also sheds light on the link between aggregates in liquid and surface attached biofilms^7^. Multiple studies have found that bacteria in aggregates have physiological characteristics that overlap with found in biofilms, including reduced antibiotic susceptibility^1–4^. This has led to the idea that aggregates can be considered as ‘non-surface attached biofilms’^1,7^, and/or as seeds for biofilm growth^5,22^. Consistent with this picture, our finding of lysis-triggered aggregation mirrors existing knowledge that biofilm formation can be triggered by the release of eDNA, caused by lysis of a subpopulation of bacteria - triggered by antibiotic, detergent, prophages or phage genes^13–15,20,23–28^. However, other aspects of our work contrast with the picture of aggregates as ‘non-surface attached biofilms’. In contrast to previous studies on aggregate physiology^1–4^, we do not observe enhanced antibiotic susceptibility of lysis-triggered aggregates compared to unaggregated cells, and we find no evidence for these aggregates acting as a precursor to biofilm formation - on the contrary, lysis-triggered aggregation appears to suppress biofilm formation. Recent work in *Pseudomonas aeruginosa* points to the existence of distinct aggregate types that form under different conditions^29^. Our work adds to this discussion, suggesting that antibiotic-induced lysis may trigger a distinct form of ‘planktonic-state’s aggregation, that happens without an associated transition to a biofilm-like physiological state.

Taken together, our work shows that low-dose antibiotic exposure can trigger the formation of *Escherichia coli* aggregates in liquid suspension. For dense, well-shaken suspensions of *E. coli*, we find that very low concentrations of the beta-lactam antibiotic mecillinam cause the formation of bacterial aggregates within hours. We link aggregation to the lysis of a small subpopulation of bacteria, releasing eDNA that mediates bacterial cohesion. In contrast to previous observations, bacteria in these aggregates appear to maintain the planktonic physiological state, since aggregation does not protect bacteria against antibiotic exposure, nor does it enhance biofilm formation. Our work contributes to increasing understanding of the nature of bacterial aggregates and the factors that can trigger their formation, and well as of the consequences of widespread low-dose antibiotic exposure in the environment and in the body.

## Results

### Low-dose *β* -lactam antibiotics cause bacterial aggregation via lysis of a few cells

To investigate the effects of subinhibitory antibiotic exposure on liquid-phase bacterial cultures, we grew cultures of *E. coli* strain MG1655 in MOPS minimal media with glucose to an optical density OD_600_ ∼0.2, corresponding to about 1.6 × 10^8^ cells/ml (see Methods), in a shaken conical flask. We then added very low doses of antibiotic, corresponding to 1/8000 times the minimum inhibitory concentration (MIC) value, measured under the same conditions (MIC^OD0.2^; see Methods and Table 1). The cultures were sampled taken over a period of 5 hours after antibiotic addition and examined using phase-contrast microscopy. At such low concentrations, the antibiotic did not significantly affect the population-level growth dynamics, as measured either by spectrophotometry or by viable counts (Supplementary Figure S1).

**Table 1.** Minimum inhibitory concentration (MIC) for mecillinam (Mec.), aztrenoam (Azt.), streptomycin (Str.), tetracycline (Tet.), and rifampicilin (Rif.) for *E. coli* strain RJA002 in MOPSGlu medium at 37 °C. MIC values were measured for cultures under different conditions: cultures at OD 0.2 at the time of antibiotic addition (MIC^OD0.2^); cultures at OD 0.8 at the time of antibiotic addition (MIC^OD0.8^); cultures that had been pre-incubated with Mec. (at MIC^OD0.2^/8000) for 4 h to induce aggregate formation (MIC^Mec.^); cultures that pre-incubated with Mec. for 4 h then treated with DNase I (MIC^Mec.*/*DNase^) to disperse the aggregates (see Methods). For details of the MIC measurement protocol, see Methods. N.M. indicates ‘not measured’. The ‘low-dose’ antibiotic concentrations used in this work and referred to in the text as MIC^OD0.2^/8000 were 1/8000 times the values reported in the top row of the table, i.e. 0.1 *μ*g/ml for mecillinam, 0.08 *μ*g/ml for aztreonam, 0.0004 *μ*g/ml for streptomycin, 0.0002 *μ*g/ml for tetracycline and 0.01 *μ*g/ml for rifampicin.

| Antibiotic | Mec. | Azt. | Str. | Tet. | Rif. |
| --- | --- | --- | --- | --- | --- |
| MIC<sup>OD0.2</sup> ($\mu\text{g/ml}$) | 840 | 640 | 3 ± 1 | 1.5 ± 0.5 | 100 ± 20 |
| MIC<sup>OD0.8</sup> ($\mu\text{g/ml}$) | N.M. | N.M. | 3 ± 1 | 2 ± 1 | 40 ± 10 |
| MIC<sup>Mec.</sup> ($\mu\text{g/ml}$) | N.M. | N.M. | 2 ± 1 | 0.4 ± 0.1 | 20 ± 10 |
| MIC<sup>Mec./DNase</sup> ($\mu\text{g/ml}$) | N.M. | N.M. | 2 ± 1 | 0.4 ± 0.1 | 20 ± 10 |

### Mecillinam and aztreonam cause morphological changes and aggregation

We tested four antibiotics - mecillinam, aztreonam, streptomycin and tetracycline. Mecillinam and aztreonam are cell-wall synthesis-targeting *β*-lactam antibiotics, while streptomycin and tetracycline bind to ribosomes, inhibiting protein synthesis^30,31^. For the two protein synthesis-targeting antibiotics, we observed no change in cell morphology, and no aggregate formation (Supplementary Figures S2 and S3). However, both the *β* -lactam antibiotics caused morphological changes, lysis of a few cells, and the formation of cell aggregates over a period of 2-4 hours (Figure 1; Supplementary Figures S4 and S5).

**Figure 1.**
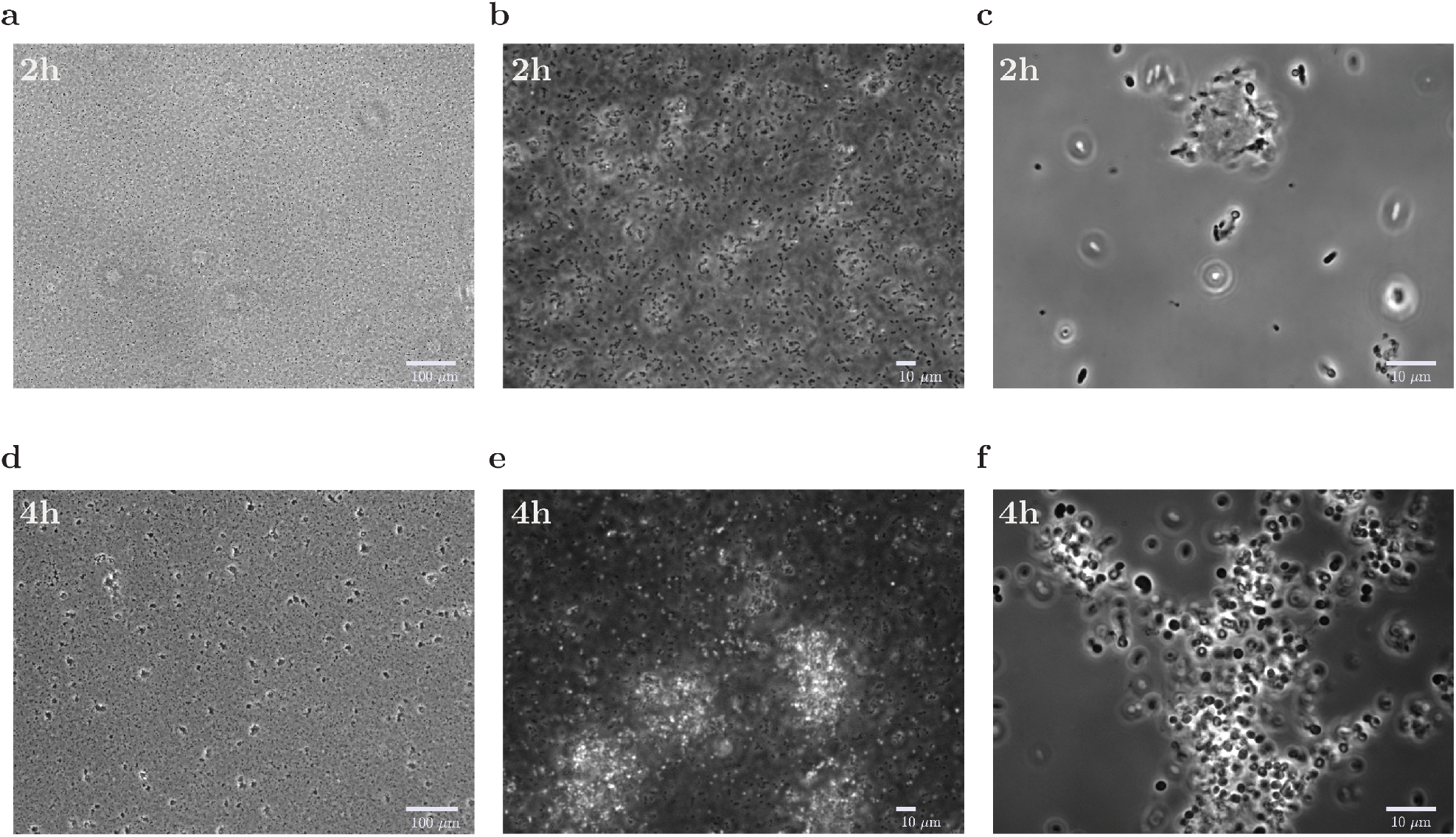
Aggregation upon incubation with mecillinam. Cultures of *E. coli* MG1655 at OD 0.2 were incubated with mecillinam at MIC^OD0.2^/8000 = 0.1*μ*g/ml, in shaken flasks (see Table 1). Phase-contrast images are shown at increasing magnification (left to right, 10× (a,d), 40× (b,e), and 100× (c,f)), for samples taken 2h (top; a,b,c) and 4h (bottom; d,e,f) after addition of mecillinam.

It is well known that *β* -lactam antibiotics cause morphological changes and lysis at high concentrations, close to or above the MIC. Mecillinam inhibits the synthesis of peptidoglycan by binding to the PBP2 transpeptidase enzyme that is required for cell elongation, causing cells to become round and lyse^32^, while aztreonam inhibits cell division by targeting the PBP3 transpeptidase, which is involved in the synthesis of cell pole peptidoglycan - hence it causes filamentation and eventually lysis^33^. Even though the concentrations of mecillinam or aztreonam in our experiments were far below the MIC, we observed similar morphological changes. For mecillinam, the cell shape changed from rod-shaped to round (Supplementary Figure S2), and swimming motility stopped, about 2 hours after addition of the antibiotic. For aztreonam, we observed filamentation (Supplementary Figure S2) and loss of motility.

For both mecillinam and aztreonam, we observed formation of aggregates consisting of 4-6 cells, about 2h after addition of antibiotic (Figure 1-b-c). After a further 30 min, these aggregates had become larger, and they continued to grow over the next 4 h (Figure 1-d-f; Supplementary Figures S4 and S5).

### Low-dose mecillinam causes a very low rate of cell lysis

Motivated by reports of biofilm formation triggered by *β* -lactam mediated lysis^13–15^, we hypothesized that the very low levels of cell lysis that we observed via microscopy (Supplementary Figure S2) might be relevant for aggregation - even though the population as a whole continues to grow under these very low levels of antibiotic (Supplementary Figure S1). To quantify the rate of antibiotic-induced cell death in our cultures, we used resazurin, a weakly fluorescent, non-toxic, cell-permeable dye which, in the presence of metabolically active cells, is irreversibly transformed to the highly fluorescent resorufin^34^. We measured resorufin fluorescence emission for exponentially growing bacterial cultures in the presence or absence of low-dose mecillinam (see Methods). Comparing the exponential rate of fluorescence increase for cultures with and without mecillinam, we concluded that about 6.4% of cell lifetimes end in lysis rather than in further division (see Methods). This low rate of cell lysis is not detectable when we compare growth curves measured via either optical density or colony forming units in the presence and absence of mecillinam (Supplementary Figure S1; see Methods).

### Aggregate formation is mediated by extracellular DNA

Extracellular DNA (eDNA) released from lysed cells has been found to play an important role in antibiotic-triggered biofilm formation e^13–15^. Therefore we speculated that released components of the lysed cells in our cultures, such as DNA or proteins, might mediate aggregation of the remaining, non-lysed cells.

### Digestion of eDNA, but not eProtein, eliminates aggregation

As a first test of whether eDNA or extracellular protein (eProtein) play a role in aggregation, we generated cultures containing aggregates by incubating with low-dose mecillinam for 5 hours (see Methods), then added either deoxyribonuclease I (DNase I), to digest eDNA, or proteinase K^2,35^, to digest eProtein (see Methods). After adding DNase I, the aggregates quickly disappeared (Supplementary Figure S4). In contrast, addition of proteinase K had no effect on the aggregates, even when we increased the concentration and incubation time (Supplementary Figure S6). Equivalent results were obtained upon adding DNase I or proteinase K to aggregates formed by incubating with low-dose aztreonam: again, pre-existing aggregates were destroyed by digestion of eDNA but not by digestion of eProtein (Supplementary Figures S5 and S6).

We also observed that DNase I could prevent aggregates from forming in the first place. Repeating our experiment, but this time adding DNase I to the dense planktonic culture at the start, together with low-dose mecillinam (or aztreonam), we observed no aggregate formation over a 5 hour period (Supplementary figure S7). In contrast, when we added proteinase K at the start of the experiment together with low-dose mecillinam, aggregates formed, suggesting that eProtein is not involved in aggregation process (Supplementary Figure S8). Repeating the same experiment with aztreonam plus proteinase K, we did not observe aggregate formation (Supplementary Figure S8), but it appeared that proteinase K might have inhibited the action of aztreonam, since we observed no change in the shape or swimming motility of the bacteria in this experiment (in contrast to our other experiments with aztreonam; Supplementary Figure S2).

These results suggest that aggregation in our experiments is mediated by eDNA, which is released when a minority of cells in the culture lyse under the action of the cell-wall targeting antibiotic.

### eDNA is present in aggregates and appears to bridge between cells

To investigate the role of eDNA in the structure of the aggregates, we performed microscopic imaging in the presence of several DNA-binding dyes. TOTO-1 is often used to stain eDNA, since it increases its fluorescence intensity by 100-to 1000-fold upon binding large fragments of DNA (10 to 50 kbp.)^36–38^. TOTO-1 binds to both dsDNA and ssDNA with similar affinity^39^; it has low penetrance of the cell membrane and therefore is not expected to stain chromosomal DNA. Propidium iodide (PI) has similar properties, but it has higher membrane penetrance and is therefore expected to stain both eDNA and chromosomal DNA of cells with damaged membranes^40^. We added TOTO-1 or PI to dense planktonic cultures, adding at the same time low-dose mecillinam or aztreonam to promote aggregation. We then imaged the cultures after 4 hours. Both dyes revealed the presence of eDNA in the aggregates (Figure 2 and Figure 3; see also Supplementary Figures S4 and S5). For both dyes, fluorescence was most intense at the positions where aggregates were observed by phase-contrast microscopy (Figure 2-d-e for TOTO-1; Figure 2-f for PI). We were even able to observe apparent eDNA ‘bridges’ between cells within the same aggregate (upper left inset of Figure 2-f).

**Figure 2.**
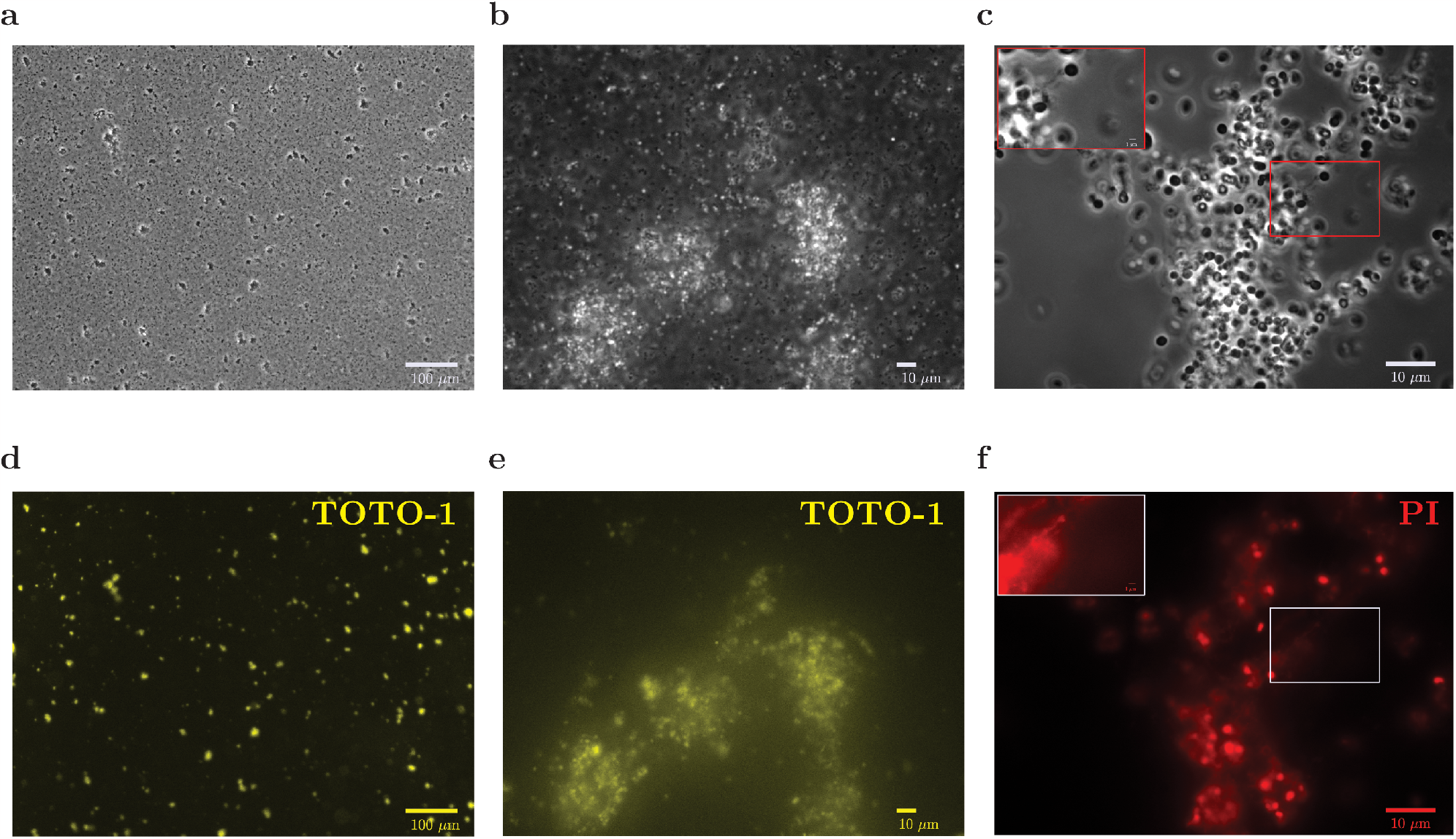
Aggregates contain eDNA. Cultures of *E. coli* MG1655 at OD 0.2 were incubated with mecillinam at MIC^OD0.2^/8000= 0.1*μ*g/ml, in shaken flasks (see Table 1). Samples were taken 4h after addition of mecillinam and stained with either TOTO-1, which binds to eDNA or propidium iodide (PI), which binds to both eDNA and the DNA of membrane-damaged-cells. The top row shows the same phase-contrast images as in Fig. 1, with different magnifications, increasing left to right: a) 10×, b) 40×, and c) 100×. The bottom row shows the corresponding fluorescence images for TOTO-1 (d,e) and PI (f). In panels (c) and (f), the upper left inset shows an expanded image of the central boxed region in which the intensity of each pixel has been increased by factor of 3.8. The inset shows the presence of eDNA bridges between cells.

**Figure 3.**
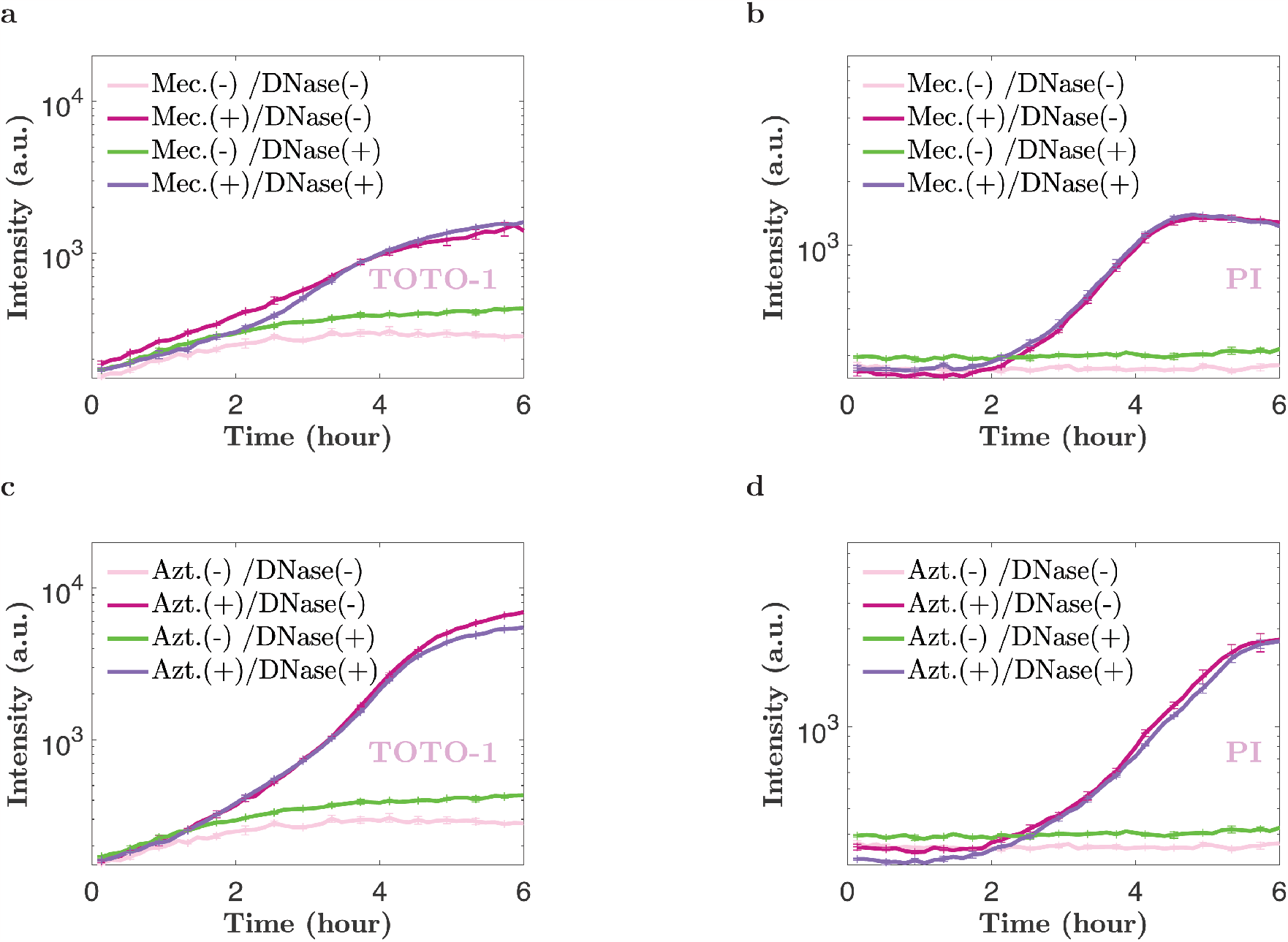
eDNA is released upon incubation with low-dose mecillinam or aztreonam. Cultures of *E. coli* MG1655 at OD 0.2 were incubated with mecillinam or aztreonam at MIC^OD0.2^/8000 (see Table 1), in shaken flasks, together with either TOTO-1, which binds to eDNA or propidium iodide (PI), which binds to both eDNA and the DNA of membrane-damaged-cells. The panels show time-series of fluorescence emission, measured using bulk spectrophotometry. TOTO-1 and PI emission was measured as a function of time (after addition of antibiotic), for cultures with and without antibiotic, and with and without DNase. Panel (a) shows results for mecillinam and TOTO-1, panel (b) shows results for mecillinam and PI, panel (c) shows results for aztreonam and TOTO-1, panel (d) shows results for aztreonam and PI. The data are expressed as mean ± standard error of the mean, for four replicate experiments.

After imaging the aggregated samples, we added DNase I to the culture and continued to take samples for imaging with TOTO-1 staining. As expected, addition of DNase I caused the aggregates to rapidly disappear in the phase contrast images (Supplementary Figure S4 for mecillinam; Supplementary Figure S5 for aztreonam); simultaneously, the TOTO-1 signal became homogeneous across the sample (Supplementary Figures S4 and S5).

### Spectrophotometry suggests eDNA release mediates aggregation

Since both TOTO-1 and PI increase their fluorescence intensity upon binding to DNA, we can also use bulk spectrophotometry measurements to detect changes in the amount of eDNA in the culture. Figure 3-a,c shows time series data for TOTO-1 emission intensity over 6 hours, following addition of low-dose mecillinam or aztreonam. The intensity of TOTO-1 emission increased by about 10-fold, 1-2 hours after addition of the cell-wall targeting antibiotic, consistent with the time at which cells were observed to change morphology and begin to aggregate. Similar results were obtained for the PI emission intensity (Figure 3-b,d), supporting the view that eDNA release mediates aggregation.

Repeating these experiments using the ribosome-targeting antibiotics streptomycin and tetracycline, for which we did not observe aggregation (Supplementary Figure S3), we observed no significant increase in emission intensity for either TOTO-1 or PI (Supplementary Figure S9), supporting our hypothesis that lysis caused by cell-wall targeting antibiotics was responsible for the eDNA release.

### Purified genomic DNA does not trigger aggregation

If aggregation is mediated by eDNA, one might expect that addition of purified genomic DNA (gDNA) would cause cells to aggregate, even in the absence of cell-wall targeting antibiotics. However, we did not observe aggregation when we added gDNA (from *E. coli* strain B at 0.25-20 *μ*g/ml final concentration) to a dense *E. coli* culture (OD_600_ = 0.2; strain MG1655). We repeated these experiments using *E. coli* strain AD37, which is unable to swim due to paralysed flagella (see Methods), to mimic the cessation of swimming motility that happens upon addition of low-dose mecillinam or aztreonam (having verified that strain AD37 forms aggregates upon addition of low-dose mecillinam and aztreonam - Supplementary Figure S10). We also varied the optical density of the AD37 cultures, testing cultures at OD_600_ = 0.2 and 0.4. However, no aggregation was observed.

This result is reminiscent of the observation of Turnbull *et al*. that exogenous addition of *P. aeruginosa* gDNA did not restore biofilm formation in a mutant that was unable to release eDNA via prophage-mediated “explosive cell lysis”^28^. Turnbull *et al*. found that, contrary to expectations, exogeneous gDNA actually inhibited microcolony formation^28^. They speculated that eDNA might need to be provided in high local concentrations to trigger biofilm initiation, or that other components might be required in addition to eDNA. Our results might suggest that other factors are required to trigger, or mediate, aggregation, in addition to eDNA. However we note also that the eDNA that is released in our experiments upon antibiotic lysis is likely to differ from purified gDNA, since DNA purification degrades DNA-associated proteins, decreases the viscosity of the DNA and may alter the composition of counter-ions on the DNA^35,41,42^.

### Ecological consequences of aggregate formation

Previous work has linked bacterial aggregation with decreased susceptibility to antibiotics^2–4^, biofilm-like bacterial physiology^1^ and the seeding of surface-attached biofilms^5^, leading to speculation that bacteria in aggregates might be in a biofilm-like physiological state, or that bacterial aggregates could be considered as a ‘third lifestyle’ ^7^. Therefore we wondered whether the bacterial aggregates that are triggered in our system by low doses of cell-wall targeting antibiotic might show altered susceptibility to other antibiotics (at much higher doses), and/or increased propensity for biofilm formation.

### Aggregation does not protect bacteria from antibiotic

To determine whether aggregate formation altered antibiotic susceptibility, we measured MIC values (MIC^Mec.^) for the antibiotics tetracycline, streptomycin and rifampicin, four hours after the addition of low-dose mecillinam to a dense culture to induce aggregation (see Methods and Table 1). Tetracycline, streptomycin and rifampicin were used because they had previously been found not to cause aggregation (Figure 3, Supplem)entary Figures S3 and S9. For comparison, we performed two control experiments. In the ‘DNase-dispersed’ control experiment, aggregates were dispersed by adding DNase I just before the MIC measurement (MIC^Mec.*/*DNase^). In the ‘no mecillinam’ control experiment we measured MIC values for equivalent cultures that had been incubated for 4 h without addition of mecillinam, reaching an OD of ∼0.8 (MIC^OD0.8^). If aggregation protects bacteria from antibiotic, then we would expect to obtain higher MIC values for the aggregated samples (MIC^Mec.^) compared to either the dispersed control (MIC^Mec.*/*DNase^) or the no mecillinam control (MIC^OD0.8^).

Comparing the MIC measurements for the aggregated samples (MIC^Mec.^) with the DNase-dispersed control (MIC^Mec.*/*DNase^) we observed no significant difference (Table 1). Therefore the presence of aggregates, in itself, has no effect on antibiotic susceptibility in our system.

However, a more complex picture emerged upon comparing the MIC values for the no-antibiotic control (MIC^OD0.8^) with those for the aggregated samples (MIC^Mec.^) or the DNase-dispersed control (MIC^Mec.*/*DNase^). For tetracycline, MIC^OD0.8^ was significantly higher than MIC^Mec.^ or MIC^Mec.*/*DNase^. This suggests that very low-dose mecillinam treatment can increase the susceptibility of *E. coli* to tetracycline (by a factor of 5), in a manner that is independent of aggregate formation. For streptomycin and rifampicin the apparent difference between MIC^Mec.*/*DNase^, MIC^Mec.^ and MIC^Mec.*/*DNase^ was not significant. Previous studies of the combined effect of tetracycline and *β* -lactam antibiotics (at much higher concentrations) suggest that these antibiotic classes tend to be antagonistic for *E. coli* cultures at low density on rich medium: i.e. the drugs are less effective when combined than they would be individually^43,44^. Therefore our result, that low-dose mecillinam can potentiate the action of tetracycline, hints that environmental conditions (imbalance in concentration between antibiotics, growth medium, cell density) might alter the nature of drug interactions.

### Biofilm formation is stimulated by low-dose antibiotic but impeded by aggregation

Low-dose antibiotic treatment has previously been found to enhance the formation of biofilms on surfaces^13–15^, while other studies have suggested that cell aggregates may be a precursor to biofilm formation^5,7,22^. To determine whether biofilm formation was affected by low-dose mecillinam treatment and/or the resulting cell aggregation, we performed microtitre plate biofilm assays. Briefly, cultures were incubated under static conditions in a 96 well microplate for 15 h, then biofilm formation was assayed using safranin O staining (see Methods and Supplementary Figure 11). To disentangle the effects of aggregation and of mecillinam treatment, we compared biofilm formation for aggregated cultures (5 h after addition of low-dose mecillinam), no-mecillinam control cultures (which had been incubated for 5 h without addition of mecillinam), DNase-dispersed control cultures (which had been incubated with mecillinam for 5 h then treated with DNase I to disperse aggregates) and vortex-dispersed control cultures (in which aggregates had been dispersed by intense mixing).

We observed a significant increase in biofilm formation for all the cultures that had been treated with low-dose mecillinam – i.e. the aggregated cultures and the two cultures in which aggregates had been dispersed (Figure 4). Therefore, low-dose mecillinam treatment led to increased biofilm formation, consistent with previous work with other cell-wall targeting antibiotics and bacterial species^14,15^. However, this increase in biofilm formation was not mediated by aggregation. Comparing our results for the aggregated cultures to the DNase-dispersed and vortex-dispersed control cultures, we found that the dispersed control cultures actually formed significantly more biofilm than the aggregated cultures (Figure 4). Therefore, aggregated cells were less able to form biofilm than dispersed cells, suggesting that for this system, aggregate formation is not a precursor to biofilm formation^5,7,22^.

**Figure 4.**
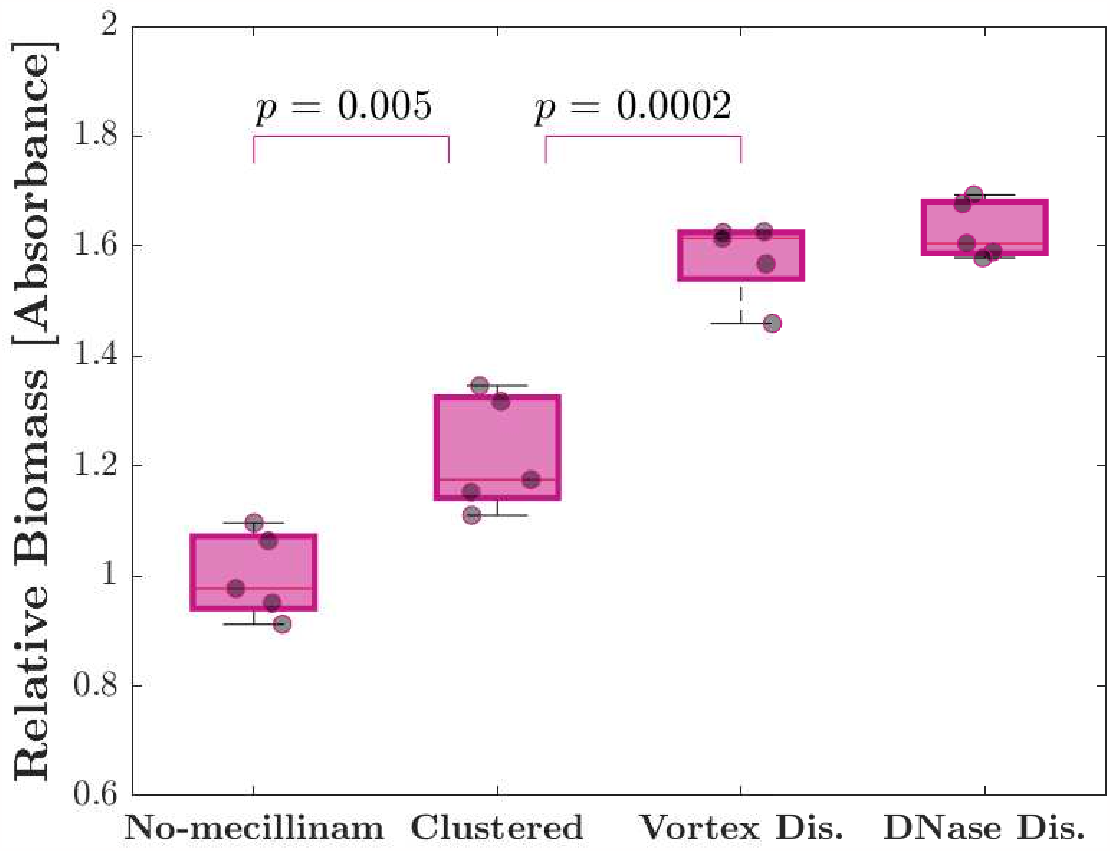
Low-dose mecillinam treatment enhances biofilm formation but aggregation does not. Relative biofilm formation was quantified in a microplate assay for (i) no-mecillinam control cultures (MG1655, MOPSGlu medium, 37 °C, initial OD 0.2, which had been incubated for 5 h without addition of mecillinam); (ii) aggregated cultures (as in (i) but incubated for 5 h with low-dose mecillinam at MIC^OD0.2^/8000); (iii) vortex-dispersed (Vortex Dis.) control cultures (as in (ii) but aggregates were mechanically disrupted); and (iv) DNase-dispersed (DNase Dis.) control cultures (as in (ii) but aggregates were chemically disrupted using DNase I) – see Methods for details. Biofilms were allowed to form for 15 h in 96-well microplates under static conditions, washed and stained with safranin O. Absorbance was measured at 510 nm and normalized to the mean absorbance of the no-mecillinam control samples. Each data point represents a single biological replicate (mean of 20-24 technical replicates). The bottom and top edges of each box indicate the 25th and 75th percentiles of five biological replicates and the red line in each box represents the median of five biological replicates. Error bars in each box are the 95% confidence interval of five biological replicates.

Our results might suggest that biofilm formation is triggered not by the release of eDNA per se, but rather by another factor that is released upon antibiotic-mediated cell lysis. An alternative hypothesis might be that released eDNA does trigger biofilm formation, but the amount of eDNA in our system may be sequestered by the aggregated cells, such that it is not available to facilitate attachment to the surface. In this scenario, dispersal of aggregates by DNase treatment or vortexing might release some eDNA, such that it can better facilitate bacterial surface attachment.

## Discussion

Microbial aggregation is ubiquitous and has significant physiological, biophysical and ecological implications, but the factors that trigger aggregation remain poorly understood. In this work, we showed that low-dose exposure to *β* -lactam antibiotics can cause *E. coli* cells to aggregate in well-shaken liquid suspension. In this system, aggregation is mediated by the release of DNA when a small subpopulation of cells lyse in response to the antibiotic. Quantitatively, we found that approximately 7% of cell lifetimes ended in lysis. While eDNA is essential for aggregation, and appears to be a key structural component of the aggregates, other factors may also be required, since purified genomic DNA did not cause aggregation in our experiments. In contrast to previous reports^2–4^, aggregation did not protect bacteria from (other) antibiotics in our system. Low-dose mecillinam treatment did promote biofilm formation, consistent with previous work^13–15^, but aggregation *per se* did not. Taken together, our results suggest a picture in which lysis of a few cells mediated by low-dose *β* -lactam treatment leads to the release of eDNA which causes aggregation, although other factors may also be involved. This is a relevant finding since suspended aggregates are widespread in industry and are increasingly being implicated in disease^2,6,7^.

### Aggregation triggered by antibiotic-mediated lysis

Although liquid phase aggregation has not been well studied, a growing body of work shows that biofilm formation on surfaces can be triggered by lysis of a subpopulation of cells and the associated release of eDNA^23–25^. This can occur through antibiotic-induced autolysis, via the activation of autolytic enzymes^14,26,27^, via the activation of prophages or upregulation of phage genes, leading to “explosive cell lysis”^13,28^, or through direct lytic action of antibiotics such as *β* -lactams^13,15^ or detergents^15,20^. Our work extends this discussion, by showing that a similar mechanism can also mediate the formation of suspended aggregates in liquid cultures.

Motivated by what is known for biofilm formation, we speculate that eDNA release in our system could occur directly through the action of antibiotic on a few susceptible cells, or could require other factors such as prophages or autolytic enyzmes. In either case, it is clear that aggregation depends critically on the presence of a small subpopulation of cells that lyse in response to the low-dose antibiotic. Stochastic bacterial response to high doses of antibiotic has been highlighted in several previous studies^45,46^ and is likely to be clinically relevant, since survival of a few cells can allow regrowth of the entire population. Our work suggests that stochastic responses to low-dose antibiotic could also have important population-level consequences. Since small populations are more susceptible to stochastic effects than large populations^47,48^, this type of aggregation might occur differently in small bacterial populations compared to large ones.

### Diverse effects of low-dose antibiotic exposure

Low concentrations of antibiotics are found in wastewater, rivers and lakes, and can also occur during antibiotic therapy, agricultural use of antibiotics or due to ecological interactions between microbes (e.g. in soil). Low-dose antibiotic exposure has been implicated in many microbiological phenomena, including enrichment of antibiotic-resistant mutants, selection for *de novo* evolution of resistance^12^, increased genotypic and phenotypic variability, and the triggering of quorum sensing and virulence^12,13^. It is also well-known that low concentrations of antibiotics can trigger biofilm formation on surfaces, not only via eDNA release as discussed above, but also via distinct mechanisms in which antibiotics act as signalling molecules, triggering gene regulatory changes associated with adhesion, metabolic stress, and expolysaccharide production^13,49^. In some cases, low-dose antibiotic exposure can also prevent biofilm formation^49^. Our work shows that the formation of liquid-phase aggregates should also be considered as a possible consequence of exposure to low doses of cell-wall targeting antibiotics. This adds to the reasons to be concerned about low-level antibiotic exposure, and strengthens the case for more extensive study of antibiotic effects at concentrations far below the MIC.

### Diversity of aggregate types

Recent work has highlighted the fact that bacterial aggregates formed under different conditions can have different structural and physiological characteristics^29^. This may help to resolve the apparent discrepancy between previous reports that liquid-state bacteria aggregates show reduced susceptibility to antibiotics^2–4^, and our observation that antibiotic susceptibility was not reduced in aggregated cells. We speculate that the aggregates formed in our experiments, which appear to be loose assemblies of cells held together by a network of eDNA strands, may retain a physiological state closer to that of planktonic cells than to that of biofilms. In contrast, dense aggregates such as those formed by *Pseudomonas aeruginosa*, seem have physiological characteristics closer to those of biofilm^1,7,50^ - although biophysical factors such as reduced antibiotic penetration into the dense aggregates may also be relevant^50^.

Insights into the diversity of aggregate types may be gained from the large body of soft matter physics work on aggregation of colloidal particles. It is well known that polymers can cause colloidal particles to aggregate via diverse mechanisms. In particular, polymer-mediated aggregation can occur either via entropic depletion interactions (which have also been observed in bacterial suspensions^4,19,51,52^), or via direct bridging of polymers between particles. Polymer bridging seems the most likely mechanism underlying the aggregation that we observe here; it has also been implicated in aggregation in other bacterial systems^53–56^.

Our system may provide a useful experimental tool to study biophysical aspects of DNA-mediated aggregation in general, since aggregation happens under well-shaken, homogeneous liquid culture conditions and the aggregates are easy to sample and image, allowing the dynamics of aggregation formation to be tracked. We also have a well-defined and controllable trigger for aggregate formation via addition of low-dose antibiotic and we can quantify the rate of antibiotic-mediated lysis. It would be productive to compare aggregation dynamics in our system with that in other aggregating liquid culture systems (e.g. *P. aeruginosa*, where aggregation is mediated by exopolysaccharides and/or eDNA under different conditions^1,3^.

### Aggregation and biofilm formation

Previous work has suggested that liquid-phase bacterial aggregates might constitute a first step on the pathway to biofilm formation^5,22^ – however we found that, although low-dose mecillinam treatment does enhance biofilm formation, this is not due to aggregation *per se*, since disruption of the aggregates by shaking or by DNase treatment leads to even better biofilm formation. This observation is consistent with our hypothesis that bacteria in these aggregates are not in a biofilm-like physiological state. To explain why aggregation seems to actually inhibit biofilm formation in our experiments, we speculate that the eDNA in our aggregates might be coated with bacteria, such that it is not available to mediate adhesion to surfaces. The size of the cloudlike aggregates that we observed is similar to the 42 *μ*m^3^ size that has been reported for *E. coli* genomic DNA in solution ^57^, suggesting that perhaps each aggregate corresponds to a single chromosome, to which bacteria have attached. These bacteria-coated chromosomes might not be good building blocks for biofilm formation. However if the aggregates are broken up by shaking or DNase, the DNA might become available for larger-scale aggregation.

### Ecological consequences of aggregation

We observed that aggregate formation via low-dose antibiotic treatment does not protect bacteria from antibiotics, nor does it enhance biofilm formation (*per se*). Since very low concentrations of antibiotics are widespread in the environment, it is tempting to speculate on what might be the ecological consequences of such aggregation. Certainly, aggregation could alter the environmental niches that are available to microbes^8^. For example, oxygen availability might become limited inside aggregates. Aggregation could also affect transport within a moving fluid. Aggregates are expected to sink faster than unaggregated cells (a crucial factor in ocean carbon dynamics as well as in wastewater treatment), and would also be transported differently in complex fluid flows such as that found in the gut - as suggested in recent work^8^. Furthermore, aggregates may interact differently with host systems such as immune cells. Interestingly, neutrophil cells can immobilise pathogens using eDNA traps (neutrophil extracellular traps, NETS^58^) – perhaps low-dose antibiotic might aid the immune system by having a similar effect. Aggregation via IgA antibodies has also been shown to prevent *Salmonella* cells from invading the gut epithelium^9^.

Although our study focused on *E. coli*, microbes are usually found in diverse communities. Within a microbiome, one would expect different species to show different aggregation responses to different stimuli – and therefore experience different biophysical or ecological consequences. This raises the interesting possibility (also hinted at in previous work^8,9^) that chemical triggers for aggregation, whether antibiotics or host factors, might provide a way to selectively control microbiome composition.

## Methods

### Bacterial Strains

The *E. coli* strains used in the project are shown in Table 2. RJA002 is a derivative of the wild-type strain MG1655 that contains a chromosomal copy of the yellow fluorescence protein gene under the control of the constitutive *λ* P_*R*_ promoter, together with a chloramphenicol resistance cassette, as previously described^59^. AD37 is a motility-deficient strain with paralyzed flagella. It contains a precise deletion of both the *motA* and *motB* genes, which were deleted simultaneously, since they are adjacent in *E. coli* MG1655. This deletion was constructed by plasmid-mediated gene replacement using the plasmid pTOF24 into which a 800 bp fragment was inserted which contained 400 bp immediately upstream and 400 bp immediately downstream of *motAB*^60^. This fragment was generated by recombinant PCR using the following four primers – Primer 1, 5’-GCAACTCGAGCCTTGAACAGTGCCCACAAGCAG-3’ Primer 2, 5’-GCAAGTCGACGCCTTTCGCTTCTAATGCCAGTT-3’ Primer 3, 5’-TGGTCAACAGTGGAAGGATGATGTCCAGCGTGAGCATGGATATAAGCGATTTTT-3’ Primer 4, 5’-AAAAATCGCTTATATCCATGCTCACGCTGGACATCATCCTTCCACTGTTGACCA-3’Restriction sites for XhoI and Sal1 were incorporated in to the flanking primers (1 and 2 respectively, sequence underlined) to permit insertion into pTOF24. This plasmid was transformed into *E. coli* MG1655 and used as a substrate for gene replacement via the 400 bp homology arms. Following selection for integration of this fragment by homologous recombination and subsequent deletion of the chromosome site, the sequence was confirmed by sequencing across the deletion using primers specific to flanking chromosomal sites.

**Table 2.** Strains used in this study.

| Strain | Parent | Characteristics | Source |
| --- | --- | --- | --- |
| MG1655 | - | K-12 wild-type | Allen lab, University of Edinburgh |
| RJA002 | MG1655 | YFP | Lloyd *et al.*<sup>59</sup> |
| AD37 | MG1655 | $\Delta motA\Delta motB$ | this study |

### Bacterial growth conditions

In the majority of our experiments, cells were grown in MOPS glucose minimal medium (MOPSGlu). This consists of potassium morpholinopropane sulfonate (MOPS) buffer, to which are added essential nutrients^61^, as well as 0.2% *w/v* glucose as a carbon source. The MOPSGlu medium was made in-house, as described by Brouwers *et al*.^62^. The doubling times of strains MG1655, RJA002, and AD37 on MOPSGlu medium at 37 °C are 60, 70, and 60± 5 min, respectively (Supplementary Figure S1).

To obtain the starting cultures for our experiments, a good-sized colony from an LB agar plate was inoculated into 5 ml MOPSGlu medium in a conical flask. The culture was then maintained in the exponential growth regime with aeration in a shaking incubator (200 r.p.m.) by diluting periodically in the same pre-warmed medium for more than 20 generations to reach a steady-state cell culture. To ensure exponential growth was maintained, sub-culturing was performed before the optical density (OD, measured at 420 nm using a Jenway 7205 spectrophotometer) reached 0.4.

### Induction of aggregation via low-dose antibiotic

To induce aggregation, planktonic bacterial cultures were prepared in steady-state, exponential phase growth via repeated serial dilution as described above. When the culture density reached 1.6× 10^8^ cells/ml (OD 0.2 @600 nm), antibiotics were added at doses corresponding to MIC^OD0.2^/8000 (with the MIC having been measured for correspondingly dense cultures on the same medium; see Table 1). The cultures were then incubated with aeration in a shaking incubator (200 r.p.m.) for periods of approximately 5 hours in a conical flask (100 ml flask containing less than 20 ml cell culture) or a falcon tube (10 ml tube containing 1-2 ml cell culture).

### MIC measurements

Our measurements of the minimal inhibitory antibiotic concentration (MIC) were performed according to previously published micro-dilution protocols^63^ with some modifications, in that we increasing the incubation time in the presence of antibiotic and we detected growth quantitatively via OD measurement, rather than with the naked eye. The detailed MIC measurement protocol is given as Supplementary Information.

### CFU measurements

The number of live cells was measured as a function of time using colony forming unit assays (CFU), for strain RJA002, during our aggregation experiments (Supplementary Figure S1). Samples for CFU measurement were taken directly from the batch culture, and the CFU per ml was measured according to the method of Herigstad *et al*.^64^. CFU measurements were performed for 3 biological replicates (independent experiments), each of which consisted of 8 replicates (technical repeats). Data were expressed as mean ± standard error of mean.

### Measuring the rate of antibiotic killing

To measure the rate of bacterial killing by low-dose exposure to mecillinam, we used resazurin sodium salt (Invitrogen), which is a weakly fluorescent, nontoxic, cell-permeable dye, that is converted to the highly fluorescent resorufin in the presence of metabolically active cells. For a bacterial population that is growing exponentially with growth rate *λ* and death rate *d*, the number of cells *N*(*t*) increases exponentially in time according to

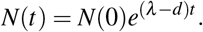

We assume that resazurin (at fixed concentration *c*_*𝓏*_) is converted to fluorescent resorufin at a rate that is proportional to the number of live (hence metabolically active) bacteria. Therefore the concentration *c* _*f*_ of resorufin increases in time according to

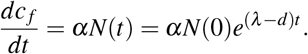

This implies that *c*_*f*_ increases exponentially according to

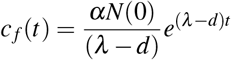

(assuming *c*_*f*_ (0) = 0). Therefore a plot of the logarithm of the resorufin fluorescence intensity as a function of time should show a straight line with gradient *m* = *λ*− *d*. For a culture without antibiotic, for which we assume no killing occurs, the equivalent plot has gradient *m*_0_ = *λ*. The antibiotic killing rate *d*, relative to the growth rate *λ*, can be found by comparing the values of *m* in the presence and absence of antibiotic: *d/λ* = (*m*_0_− *m*)*/m*. Low-dose mecillinam (MIC^OD0.2^/8000; see Table 1) was added to planktonic cultures of *E. coli* strain MG1665 grown at 37 °C in MOPSGlu medium, at optical density OD_600_ = 0.2 (∼1.6 × 10^8^ cells/ml as described above). The culture was then transferred into a 96-well black/clear flat-bottomed microplate (Falkon), such that the total volume of cell culture per well was 200 *μ*l. Resazurin was added to each well to a final concentration of 0.044 mM. The plate was incubated for 6 h at 37 °C, using double orbital shaking at 600 r.p.m. in a plate reader (CLARIOstar, BMG), covered with a Breathe-Easy sealing membrane (Sigma-Aldrich). Time series of the fluorescence emission of resorufin were recorded using a 530-560/580-600-Exc./Emi. filter at 12 min intervals. The logarithm of fluorescence intensity was plotted as a function of time and the gradient was obtained for early times (in the interval 48 min to 96 min after adding mecillinam). The experiments (Fig. 1-d in the SI file) were repeated with five biological replicates (independent experiments), and each biological replicate was repeated with three technical repeats. Data are expressed as mean ± standard error of mean. Using this procedure, the gradients that we obtained for the plots of logarithm of fluoresence signal vs time were *m*_0_ = 1.46 ± 0.04/h (in the absence of mecillinam) and *m* = 1.36 ± 0.03/h (in the presence of mecillinam). Therefore, we conclude that the killing rate is *m*_0_ − *m* = 0.1 ± 0.07/h, or equivalently, 6.8 ± 1% of bacterial lifetimes (doubling time about 60 min; MOPSGlu medium; 37 °C) end in lysis in antibiotic-mediated killing. This killing rate is too low to be detected from growth curves measured using optical density or colony-forming units (Supplementary Figure S1).

### Digestion of eDNA and eProtein in pre-formed aggregates

Aggregated samples of *E. coli* strain MG1665 were created by adding low-dose antibiotic to planktonic cultures grown in MOPSGlu medium at 37 °C, and incubating for 4 hours (as described above). Samples were then taken from the cell culture flasks. Deoxyibonuclease I (DNase I) solution (Stem Cell Technology, 1 mg/ml), and Proteinase K solution (Fisher, 500 *μ*g/ml) were used at a range of concentrations, for eDNA digestion and eProtein digestion, respectively. The addition of DNase I at 5 % *v/v* (DNase I solution: 100 *μ*g/ml) followed by incubation for 10 min completely removed bacterial aggregates generated by mecillinam addition (Supplementary Figure S4), while DNase I at 10% *v/v* (DNase I solution: 1 mg/ml) followed by incubation for 10 min removed aggregates generated by aztreonam addition (Supplementary Figure S5). We only performed eProtein digestion experiments on aggregates that were generated by adding mecillinam. We added Proteinase K solution to the samples, at final concentrations of 10 %, 20 %, or 50 % *v/v*, followed by incubation for 15 min, 30 min, or 45 min at 37 °C (all times with all concentrations). None of these proteinase K treatments had any detectable effect on the aggregates, as observed by microscopy (Supplementary Figure S6).

### Prevention of aggregation by early addition of DNase I

To test whether aggregation was prevented by addition of DNase I at the start of the experiment, cultures of strain MG1655 were prepared in steady-state exponential growth (see above), at OD_600_ = 0.2. Mecillinam or aztreonam was added at low dose (MIC^OD0.2^/8000; see Table 1), in addition to DNase I solution at 5 % *v/v* (100 *μ*g/ml) for the mecillinam experiments, or 10% *v/v* (1 mg/ml) for the aztreonam experiments. The cultures were then followed both via microscopy and via spectroscopy (using a plate reader). For microscopy, the cultures were incubated with shaking (37 °C, 200 r.p.m.) for periods of approximately 5 hours in a conical flask. Samples were removed at regular intervals and imaged under the microscope to look for aggregates (see below for imaging protocol). For spectroscopy, the cultures were transferred to a polystyrene microplate with clear flat-bottomed wells (Greiner Bio-One), such that the total volume of cell culture per well was 200 *μ*l. The microplate was incubated in a plate reader (Fluostar Optima) at 37 °C, with double orbital shaking at 600 r.p.m., for 5 h. OD_600_ was recorded every 6 minutes (Supplementary Figure S7). The experiments were performed with four technical repeats, and data are expressed as mean ± standard error of mean.

### Testing whether purified *E. coli* genomic DNA promotes aggregation

To test whether addition of purified genomic DNA promotes aggregation, in the absence of antibiotic treatment, we prepared steady-state, exponential cultures of *E. coli* strains MG1665 (motile) and AD37 (non motile) as described above, on MOPSGlu medium at 37 °C. When the cultures reached OD_600_=0.2 or 0.4, deoxyribonucleic acid sodium salt from *E. coli* strain B-Type VIII (Sigma-Aldrich), was added at 0.25 *μ*g/ml, 1 *μ*g/ml, 5 *μ*g/ml, 10 *μ*g/ml, and 20 *μ*g/ml final concentrations. The cultures were then incubated with aeration in a shaking incubator (200 r.p.m.) for up to 1.5 hours at 37 °C. The samples were imaged (see below for imaging protocol) to assess whether or not aggregates had formed. No aggregates were observed under any of the conditions tested (varying initial OD, concentration of added DNA, strain MG1655 vs AD37).

### Microscopy

Image acquisition was performed using an inverted epifluorescence microscope (Ti-U, Nikon) with 10×, 40×, 60×, and 100× (1.45 NA phase oil) objectives in combination with a digital-camera (CoolSNAPHQ2, Photometrics). Sample preparation followed different protocols depending on the magnification of the objective. To image with the 100× objective, 5 *μ*l of cell culture was sandwiched between a cover glass and a glass microscope slide, and sealed with VALAP (an equal mixture of Vaseline, Lanolin, and Paraffin). To image at lower magnifications, 15-20 *μ*l of cell culture was sandwiched between a glass cover slip and a glass microscope slide, using a 1× 1 cm^2^ disposable, double-sided adhesive plastic Gene Frame (Thermo Scientific). Once prepared, the chamber was allowed to stand for a few minutes with the cover glass at the bottom, to allow aggregates to settle onto the cover glass. In all cases, multiple images (1392 ×1040 pixels, 16 bit, 1×1 binning) were taken for different parts of the chamber.

In this work, we used two DNA stains: TOTO-1 iodide (TOTO-1, Thermo Fisher) and propidium iodide (PI, Thermo Fisher). The final concentration of each dye was 1 % *v/v*, which corresponds to 100 *μ*M for TOTO-1, and 100 *μ*g/ml for PI. Prior to imaging, dye solution was added to the cell culture, and return to the incubator to continue shaking as before. After about 5 min, sampling was done for microscopy. The fluorescence emission of TOTO-1 and PI were recorded using the microscope filters ET-EYFP (Chroma, 49003), and ET-DSRed (Chroma, 49005), respectively, and using the same time interval for excitation/emission for the same dye in all samples.

### Spectrophotometry

To detect eDNA via spectrophotometry (Fig. 3), steady-state, exponential cultures were prepared as described above, at OD_600_=0.2. The cultures were then transferred to a 96-well black/clear flat-bottomed microplate (Falkon), at a total volume of 200 *μ*l of culture per well. Low-dose antibiotic, PI, TOTO-1, and DNase I were added to each well, to final concentrations of MIC^OD0.2^/8000 for antibiotic and 1 % *v/v* for the dyes. The plate was incubated for 5 h at 37 °C, using double orbital shaking at 600 r.p.m. in a plate reader (CLARIOstar, BMG), with the lid on. Time series of the fluorescence emission of PI and TOTO-1 were recorded using a 544/590-Exc./Emi. filter and a 500/520-Exc./Emi. filter, respectively, at 10 min intervals.

The experiments were performed in three biological replicates (independent experiments), and the data are expressed as mean ± standard error of mean, while each biological replicate was performed with three technical repeats.

### Biofilm formation assay

We quantified biofilm formation for (i) unaggregated dense cultures, as a control, (ii) aggregated cultures, that had been treated with low-dose mecillinam, (iii) cultures in which aggregates were mechanically disrupted, and (iv) aggregated cultures in which aggregates were chemically disrupted using DNase I (Figure 4).

This quantification was performed using a modified version of the standard staining protocol for biofilm biomass quantification^65^ (see the detailed description in the Supplementary Information). Four 50 ml falcon tubes were prepared containing 5 ml of steady-state cell culture at OD_600_=0.2 and low-dose mecillinam (MIC^OD0.2^/8000; see Table 1) was added to 3 of them. The falcon tubes were incubated with aeration in a shaking incubator (200 r.p.m.) at 37 °C for 5 hours, after which the presence of aggregates in the samples with mecillinam (and absence of aggregates in the sample without mecillinam) was checked using optical microscopy (20× objective).

For mechanical aggregate disruption, one of the aggregated cultures was transferred to a reagent reservoir (VistaLab Technologies) and mixed 50 times with an 8-channel micropipette (Eppendorf Xplorer, with 350 *μ*l tips, taking up 200 *μ*l per channel at speed setting # 8). During this procedure care was taken to mix the entire culture by moving the pipette around the reservoir. For chemical aggregate disruption, DNase I solution at 5 % *v/v* (100 *μ*g/ml) was added to another of the aggregated cultures and mixed for 30 s (40 Hz; Fisher Scientific mixer). The culture was then transferred to a reagent reservoir and mixed 5 times with an 8-channel electronic pipette, as above. For both mechanical and chemical disruption, the disappearance of aggregates was checked using microscopy (20× objective). After the cultures had been transferred to a microplate (see below), an 8-channel pipette was used to mix (30 times) the contents of each well to ensure all aggregates were removed.

Biofilm formation was quantified using Safranin-O staining. A detailed protocol is given as Supplementary Information.

### Statistical Analysis

All statistical analyses were performed by using Matlab R2019a (MathWorks; Natick, MA, USA). One-way ANOVA was applied and the results with *p <*0.05 were considered statistically significant.

## Data and code availability

The datasets and original code generated during the current study are available from the corresponding author on reasonable request.

## Author contributions

ST and RJA designed the project. AD created and tested strain AD37 which is used in the study. ST collected and analysed the data. ST and RJA interpreted the data. ST wrote the manuscript. ST and RJA edited the manuscript.

## Acknowledgements

We thank Omar Shabana for help with the spectrometry experiments, and Sedigheh Abedi, Rebecca Brouwers, Anne Busch, Gavin Melaugh, Elin Lilja, Vijay Srinivasan and Stefanie Wagner for useful discussions and advice. We also thank Patricia Gonzalez for lab management. This work was supported by the European Research Council under Consolidator grant 682237 “EVOSTRUC”, by BBSRC under Grant No.BB/R012415/1. This work was also funded by the Deutsche Forschungsgemeinschaft (DFG, German Research Foundation) under Germany’s Excellence Strategy-EXC 2051-Project ID 390713860. For the purpose of open access, the author has applied a Creative Commons Attribution (CC BY) licence to any Author Accepted Manuscript version arising from this submission.

