## Supplemental methods and figures for "Bacterial aggregation triggered by low-level antibiotic-mediated lysis"

### Bacterial aggregation triggered by low-level antibiotic-mediated lysis: Supplementary Methods and Figures

#### Supplementary Methods

##### MIC measurement protocol

Our MIC measurement protocol was as follows. The culture was transferred into a polystyrene microplate with clear, flat-bottomed wells (Greiner Bio-One) and covered with a lid. The total volume of cell culture supplemented with antibiotic in each well was 200 $\mu$l, while care was taken to keep the maximum volume of the antibiotic solution less than 10% of the total volume for all antibiotics except for rifampicin, where the maximum volume of antibiotic solution was less than 20% of the total volume. Antibiotic was added to the wells in serial dilutions (two fold or less), with at least four replicates for each dilution. The concentration ranges for the antibiotics were as follows – mecillinam 1.64 mg/ml to 13 $\mu$g/ml; aztreonam 128 $\mu$g/ml to 10 $\mu$g/ml; streptomycin 8 $\mu$g/ml to 1 $\mu$g/ml; tetracycline 140 $\mu$g/ml to 100 ng/ml; and rifampicin 240 $\mu$g/ml to 10 $\mu$g/ml. Following antibiotic addition, bacterial growth was monitored by incubating the microplate in a plate reader (Fluostar Optima) at 37 °C, with double orbital shaking at 600 r.p.m., for 24 h (about 24 generations in MOPSGlu medium). The OD of each well was recorded every 6 minutes at 600 nm. We defined the MIC as the lowest concentration of antibiotic that prevents an increase in the OD (at the end of the experiment, *i.e.* after 24 h) greater than 5-10% of the starting OD measurement. All of our MIC measurements were performed with three biological replicates (independent experiments), and the final data were expressed as mean $\pm$ standard error of mean.

MIC measurements were performed for strain RJA002 at 37 °C in MOPSGlu medium, under various conditions. In particular, we varied the starting OD of the culture, and in some experiments we used aggregated cultures, which had been subjected to 4 h of low-dose mecillinam treatment (Table 1 of the main text). We also compared MIC values for mecillinam for all the strains used in this work (for an inoculum OD of 0.2 (at 600 nm) in MOPSGlu medium at 37 °C). We observed no significant differences in MIC between the strains.

##### Biofilm quantification protocol

To perform the biofilm assay, all 4 cell cultures were transferred into 96-well clear U-bottomed polystyrene microplates (Thermo Scientific); 200 $\mu$l per well. The samples were arranged in the plates as indicated in Supplementary Figure S11. Each sample type (i)-(iv) as defined in the Methods section of the main text occupied 20-24 wells (technical replicates). The wells in the corners of the plate were filled with sterilized water to minimize liquid evaporation. One column (6 wells) was left empty to assess the lower detection limit (LDL). Another column (6 wells) was filled with medium only for background subtraction. Then, the microplate was incubated with a lid on for 15 h, at 37 °C without shaking, to allow biofilm to form.

To remove planktonic cells and stain the biofilm, we used the following procedure. For all wells containing liquid, 100 $\mu$l of liquid was removed gently from each well with the 8-channel electronic pipette (100$\mu$l tips) at the lowest speed setting (speed # 1). To remove the rest of the liquid, each well - apart from the empty column - was washed 3 times with 200 $\mu$l of sterile phosphate-buffered saline (PBS) (using the 8-channel electronic pipette; 350 $\mu$l tip, at the lowest speed setting). The microplate was then left under a laminar hood for 1 hour to partially dry out. To stain the biofilm biomass that was adhering to the bottom of the wells, all wells (including the empty wells) were incubated with 10 $\mu$l aqueous safranin O dye (0.1 % $v/v$ in water, Scientific Laboratory Supplies) for 30 min at room temperature in the dark. The safranin O solution was then removed with the 8-channel electronic pipette (100 $\mu$l tip) at the lowest speed setting, and non-bonded stain was removed by washing 3 times with 200 $\mu$l of PBS (8-channel electronic pipette; 350 $\mu$l tip; lowest speed). The plate was then dried in an incubator at 37 °C for about

1 h (Supplementary Figure S11).

To measure the remaining biofilm biomass, the bonded stain was extracted and quantified. 200 $\mu$l of HCl (0.1 M) was added to each well and mixed with the 8-channel electronic pipette. The well contents were allowed to settle for 15 min (to allow biomass to sink) then 100 $\mu$l of liquid was removed gently with the 8-channel electronic pipette (100 $\mu$l tip, at the lowest speed setting) and transferred to a new flat-bottom microplate (Greiner bio-one). To quantify safranin O the absorbance at 510 nm (CLARIOstar, BMG) was measured for all samples (see Supplementary Figure S11 for the spectrum of safranin-O in HCl).

To account for the binding of safranin O to the plate material, for each experiment we considered the average absorbance of the empty wells in the plate as a background-staining-signal (BSS). The BSS is expressed as a mean $\pm$ standard deviations from the 6 empty wells (technical replicates). The lower detection limit (LDL) for a given experiment was then defined as the mean value of the BSS plus 3 standard deviations. Any measured values for other wells in the same experiment that were smaller than the LDL were disregarded. Furthermore, the average absorbance of the wells in the medium-only control column was used as a background measurement (mean background, MB). The MB is expressed as a mean $\pm$ standard deviations for the 6 medium-only wells (technical replicates). For all sample wells whose absorbance was higher than the LDL, the MD value was subtracted. The experiments were performed in five biological replicates (independent experiments), and the final data were expressed as mean $\pm$ standard deviations from the five biological replicates.

Finally, to compare the data to the no-mecillinam control (sample type (i) as defined in the Methods section of the main text), the average, background-subtracted absorbance for each sample type (ii)-(iv) was divided by the average absorbance for the no-mecillinam control (sample type (i)) for each biological replicate (see Figure 4 of the main text).

#### Supplementary Figures

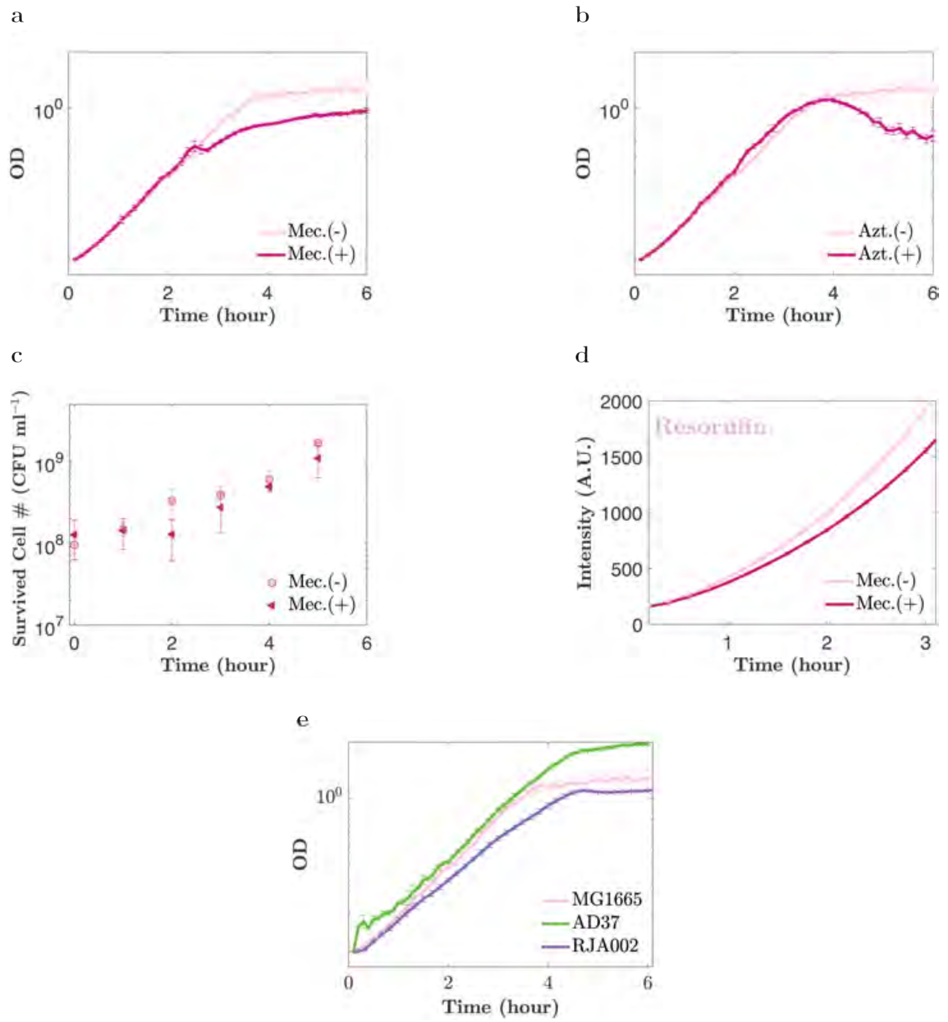


**Supplementary Figure 1.** **Low-dose antibiotic treatment has little effect on early-time population growth.** (a,b): Growth curves (optical density vs time) for the wild-type strain MG1655 on MOPSGlu media at 37 °C measured in a 96-well plate reader in the presence or absence of (a) mecillinam and (b) aztreonam, at $MIC^{OD0.2}/8000$ (see Table 1 of the main text). (c): Plate counts (CFU/ml) for samples taken from shake-flask cultures of RJA002, on MOPSGlu media at 37 °C in the presence and absence of mecillinam at $MIC^{OD0.2}/8000$. (d): Fluorescence intensity of resorufin dye, for 96-well plate cultures of MG1655 on MOPSGlu media at 37 °C in the presence and absence of mecillinam at $MIC^{OD0.2}/8000$. (e): Plate reader growth curves (OD vs time) for the strains used in this study. In all panels, data are expressed as mean ± standard error of mean, for three replicate experiments.

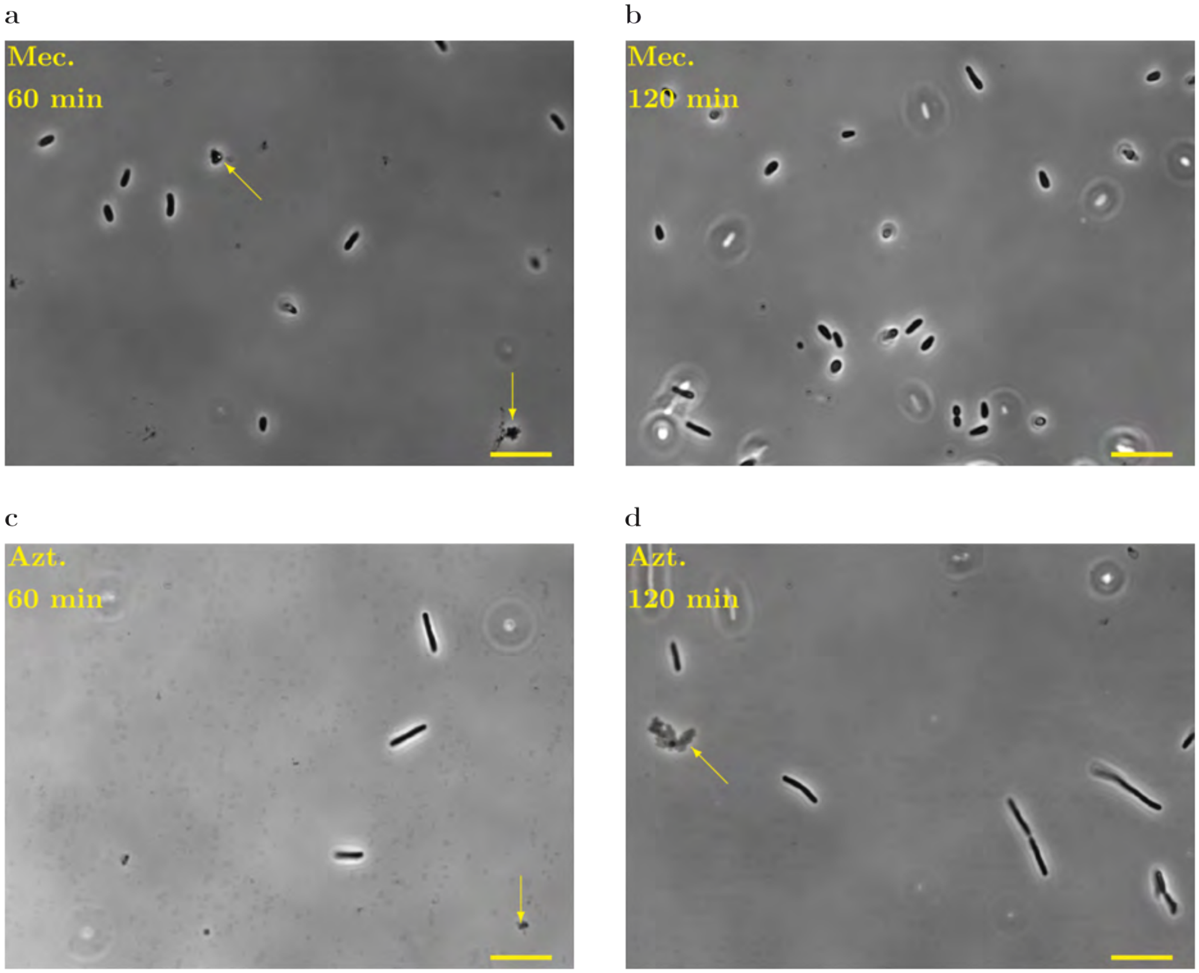


**Supplementary Figure 2. Cell morphology is affected by low doses of mecillinam or aztreonam.** *E. coli* wild-type cells (MG1655) were imaged 60 min and 120 min after addition of antibiotic at $\text{MIC}^{\text{OD}0.2}$/8000 (see Table 1 of the main text), on MOPSGlu media at 37 °C. (a,b): Low-dose mecillinam causes cells to become round. (c,d): Low-dose aztreonam causes cells to filament. A few lysed cells were visible, indicated by the arrows. Scale-bar = 10$\mu$m.

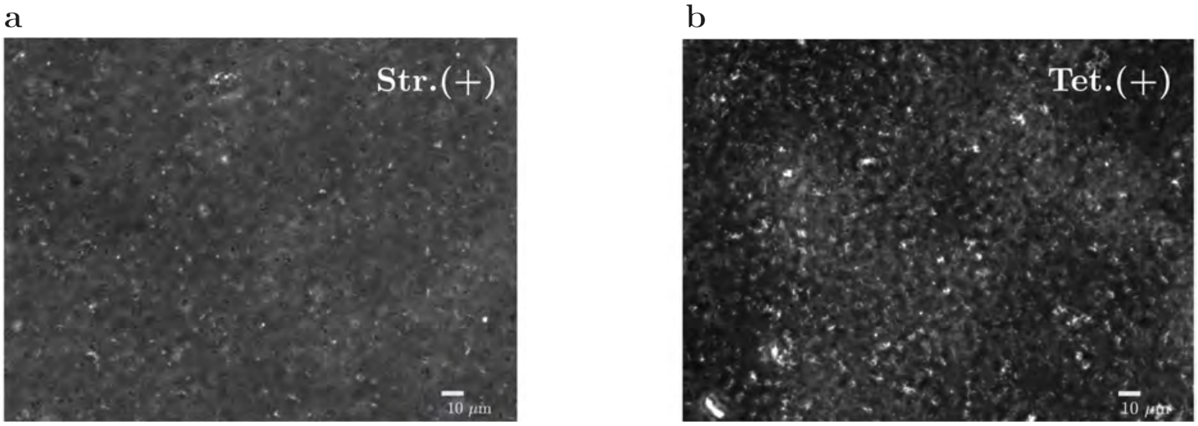


**Supplementary Figure 3. Low doses of streptomycin or tetracycline do not cause aggregation.**
Phase-contrast images (a,b), for cultures of *E. coli* MG1655 at OD 0.2 on MOPSGlu media at 37 °C incubated with streptomycin (a) or tetracycline (b) at $MIC^{OD0.2}/8000$ (see Table 1 of the main text).

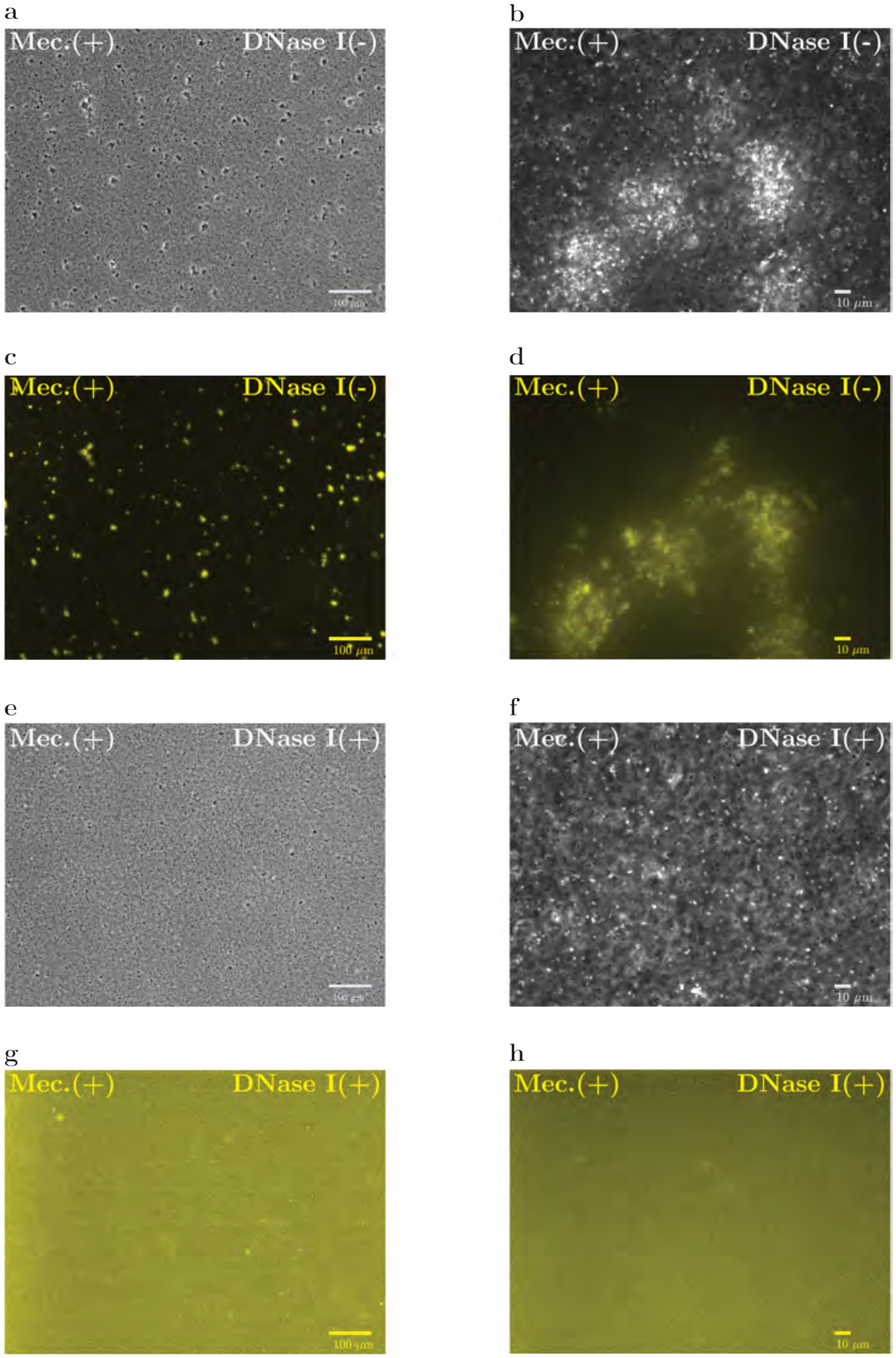


**Supplementary Figure 4. Mecillinam-triggered aggregates contain eDNA, and DNAse I treatment disperses pre-formed aggregates.** Phase contrast images and the corresponding TOTO-1 fluorescence images, for the wild-type MG1665 strain, incubated with mecillinam at $MIC^{OD0.2}/8000$, in shaken flasks (see Table 1 of the main text), for 4 h. In images labelled DNAse I(+), DNase I (5% $v/v$) was added at the end of the experiment (see Methods section of the main text).

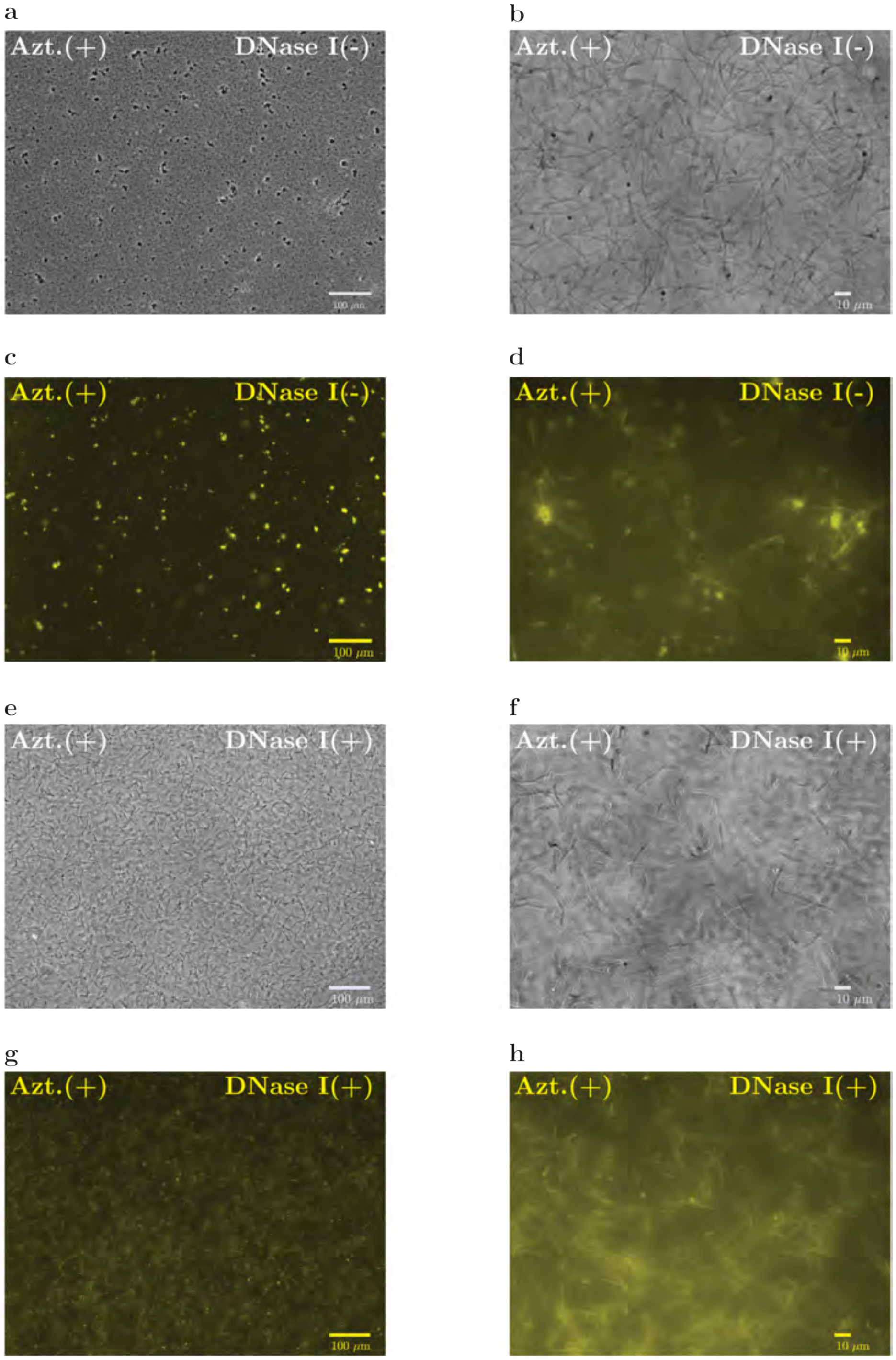


**Supplementary Figure 5. Aztreonam-triggered aggregates contain eDNA, and DNAse I treatment disperses pre-formed aggregates.** Phase contrast images and the corresponding TOTO-1 fluorescence images, for the wild-type MG1665 strain, incubated with aztreonam at $MIC^{OD0.2}/8000$, in shaken flasks (see Table 1), for 4 h. In images labelled DNAse I(+), DNase I (10% $v/v$) was added at the end of the experiment (see Methods section of the main text).

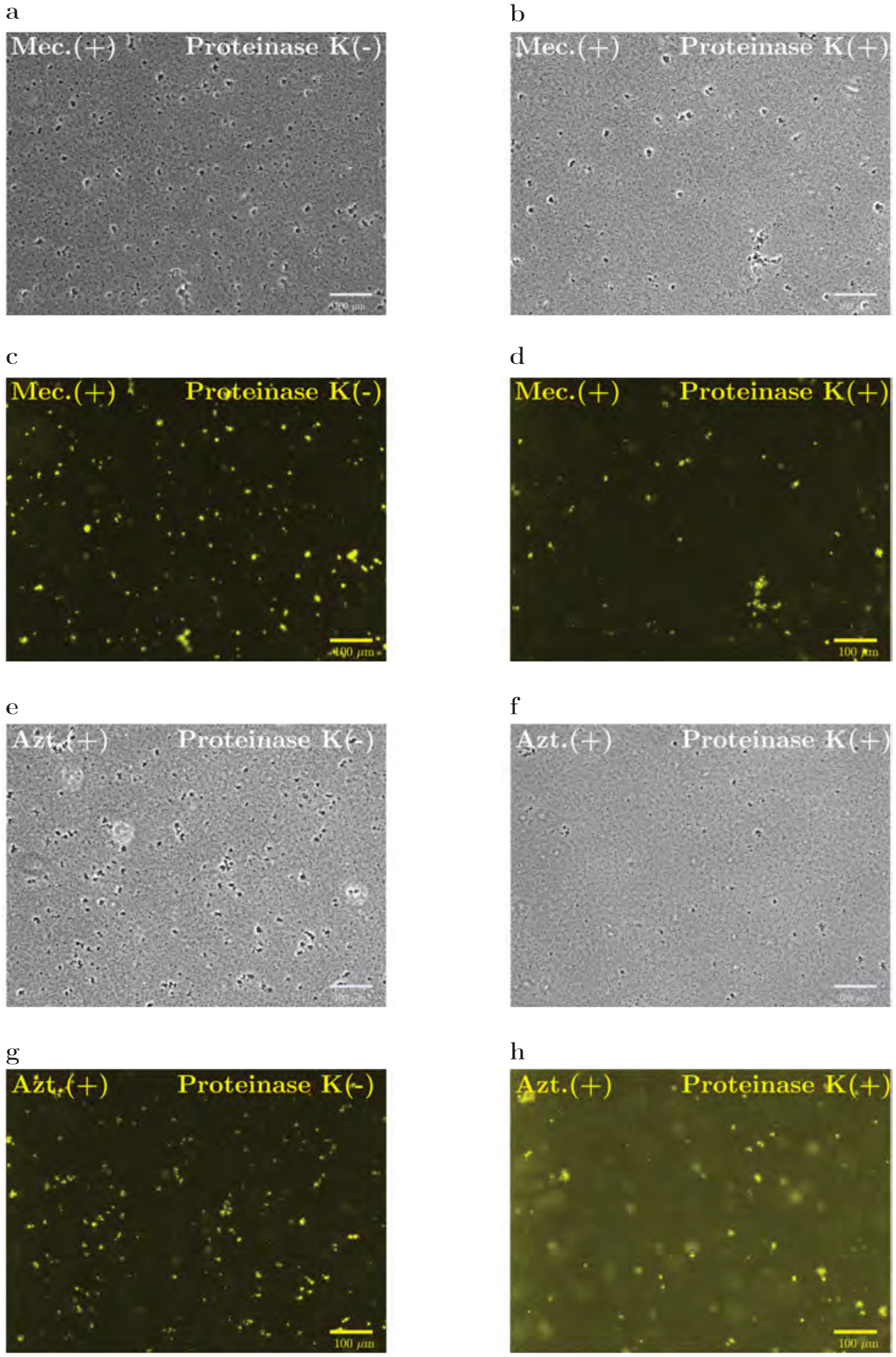


**Supplementary Figure 6. Addition of proteinase K does not destroy aggregates.** Phase contrast images and the corresponding TOTO-1 fluorescence images, for the wild-type MG1665 strain, incubated with (a-d) mecillinam or (e-h) aztreonam at $\mathrm{MIC}^{\mathrm{OD}0.2}/8000$, in shaken flasks (see Table 1 of the main text), for 4 h, with and without addition of Proteinase K (20% $v/v$), at the end of the experiment.

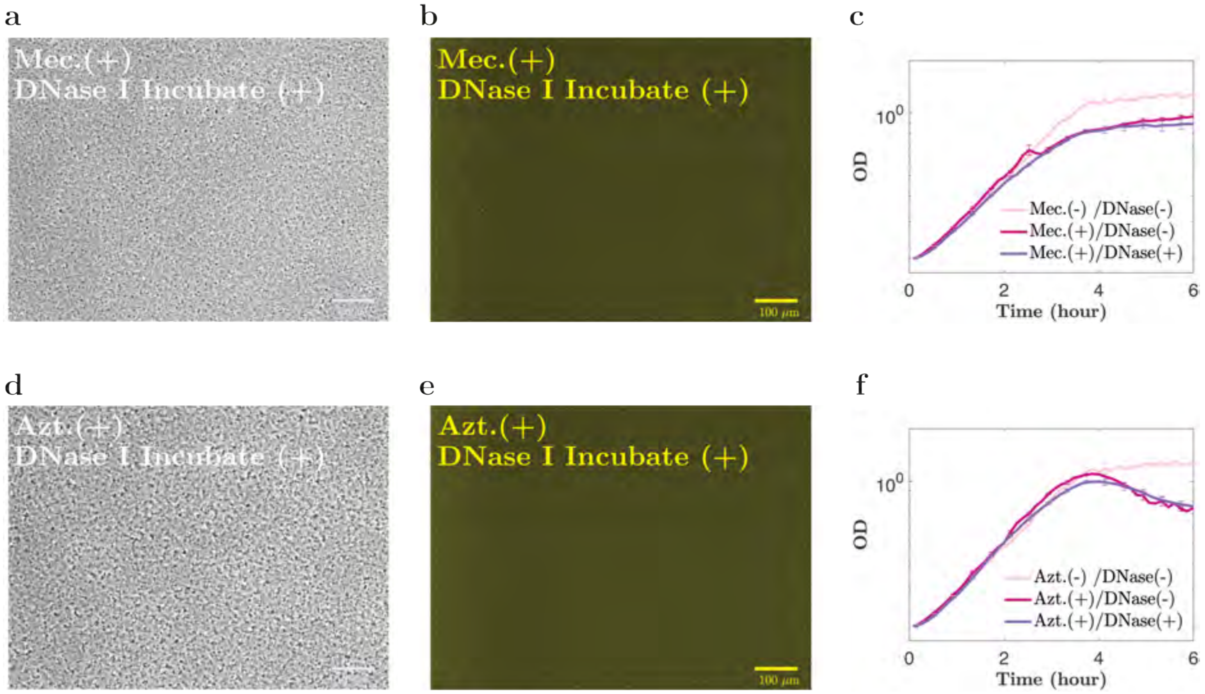


**Supplementary Figure 7. Incubation with DNAse I prevents aggregation.** (a,b,d,e): Phase contrast images and the corresponding TOTO-1 fluorescence images, for the wild-type MG1665 strain, incubated with mecillinam or aztreonam at $MIC^{OD0.2}/8000$, in shaken flasks (see Table 1), in the presence of absence of DNase I (5% $v/v$, from the start of the experiment), for 4 h. (c,f): Growth curves (OD vs time) for the same cell cultures, with and without mecillinam and aztreonam, and/or DNase I. Low-dose antibiotic treatment has no significant effect on early-time population-level growth but the antibiotic-treated cultures do show a reduction in the measured OD at later times, probably due to aggregation.

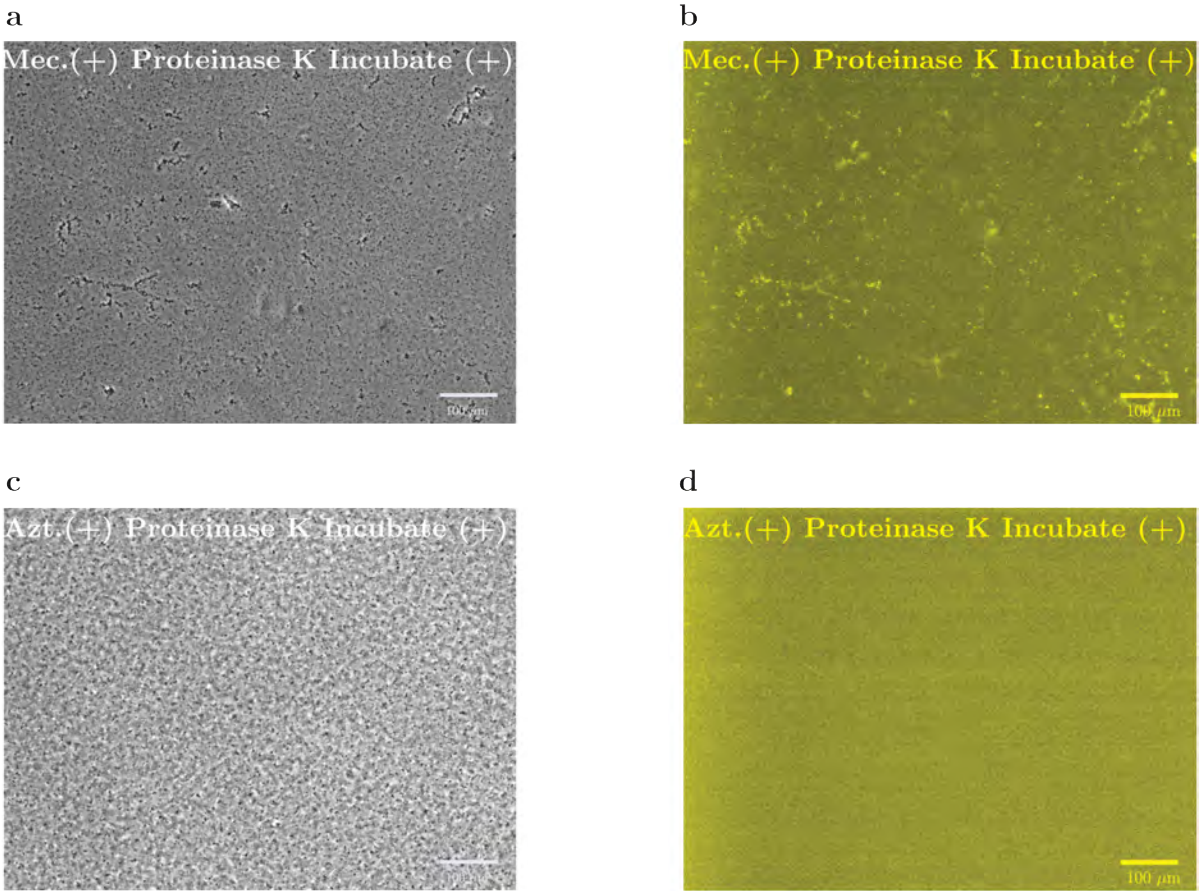


**Supplementary Figure 8. Incubation with proteinase K does not prevent aggregation.** Phase contrast images and the corresponding TOTO-1 fluorescence images, for the MG1665 wild-type strain, incubated with mecillinam or aztreonam at $MIC^{OD0.2}/8000$, in shaken flasks (see Table 1 of the main text), in the presence of proteinase K (20% $v/v$, from the start of the experiment), for 4 h.

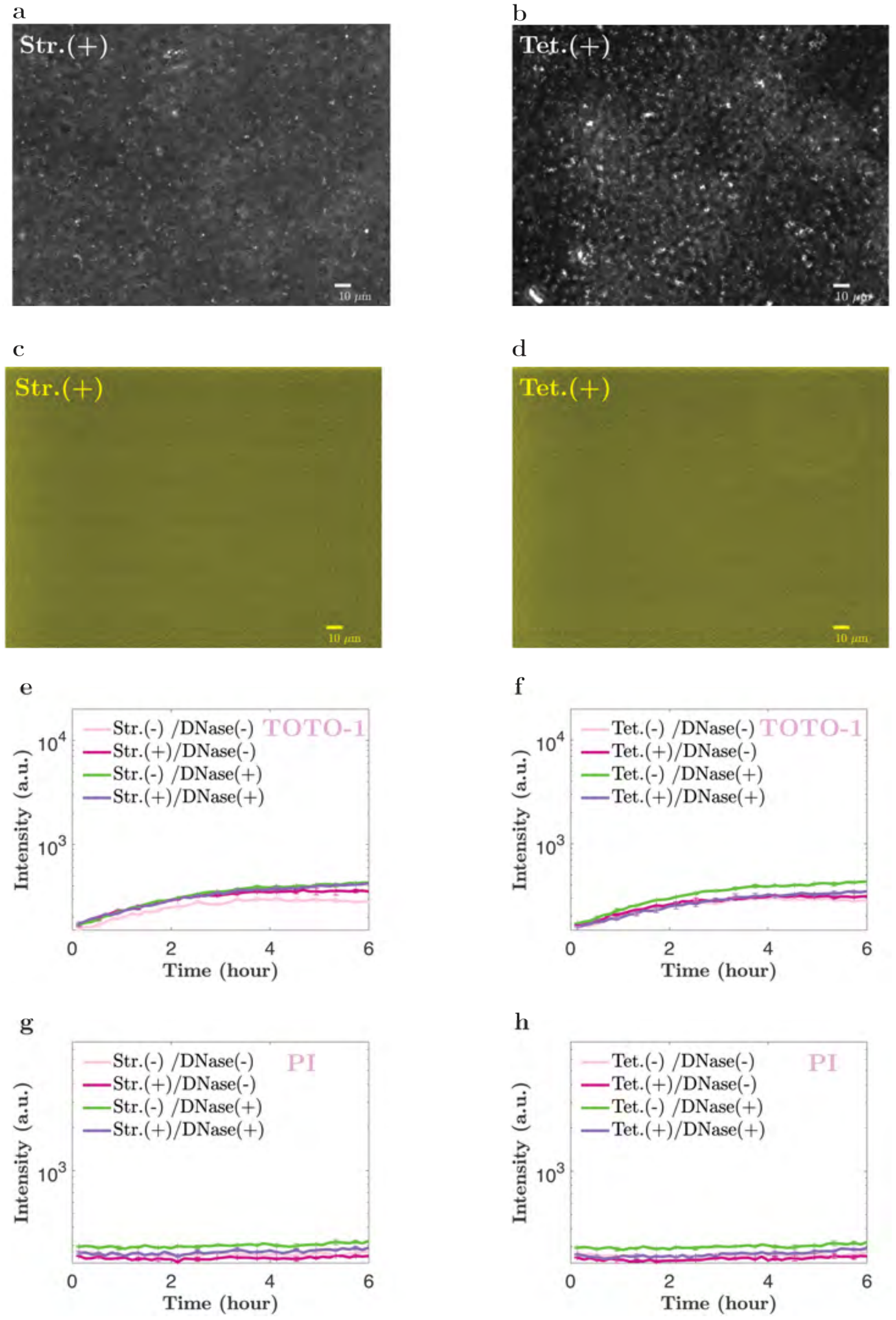


**Supplementary Figure 9. Low doses of streptomycin or tetracycline do not cause release of eDNA or aggregation.** Phase-contrast images (a,b), and the corresponding fluorescence images (c,d), for cultures of *E. coli* MG1655 at OD 0.2 on MOPSGlu media at 37 °C incubated with streptomycin (a,c) or tetracycline (b,d) at $MIC^{OD0.2}/8000$ (see Table 1 of the main text). (e-h): Bulk spectrophotometry measurements of TOTO-1 or PI fluorescence, for plate-reader cultures incubated with or without low-dose antibiotic and in the presence or absence of DNAse I. None of the cultures show increased fluorescence corresponding to eDNA release. The data are expressed as mean ± standard error of mean, for four replicate experiments.

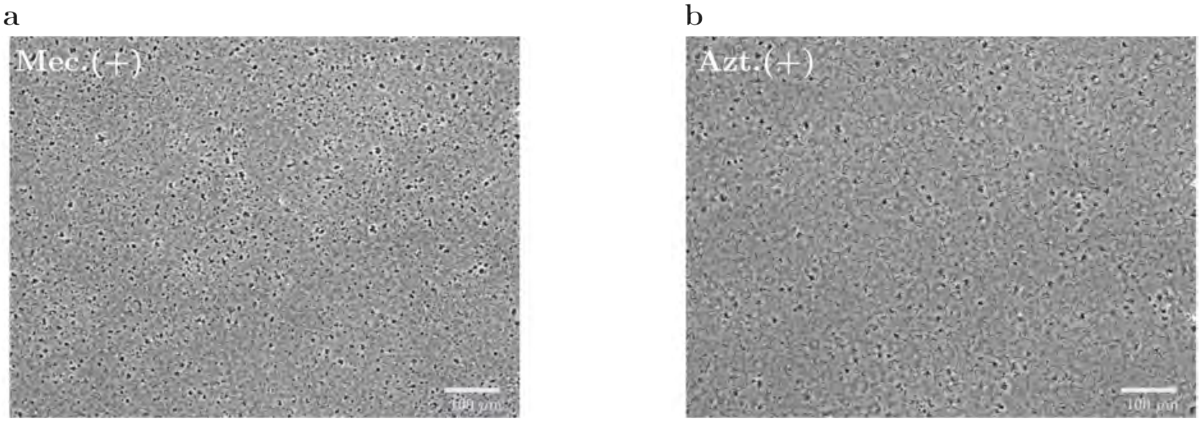


**Supplementary Figure 10. Strain AD37, which has paralysed flagella, aggregates when treated with low-dose mecillinam or aztrenam.** Phase contrast images for strain AD37, incubated with mecillinam or aztreonam at $MIC^{OD0.2}$/8000, in shaken flasks (see Table 1 of the main text), for 4 h.

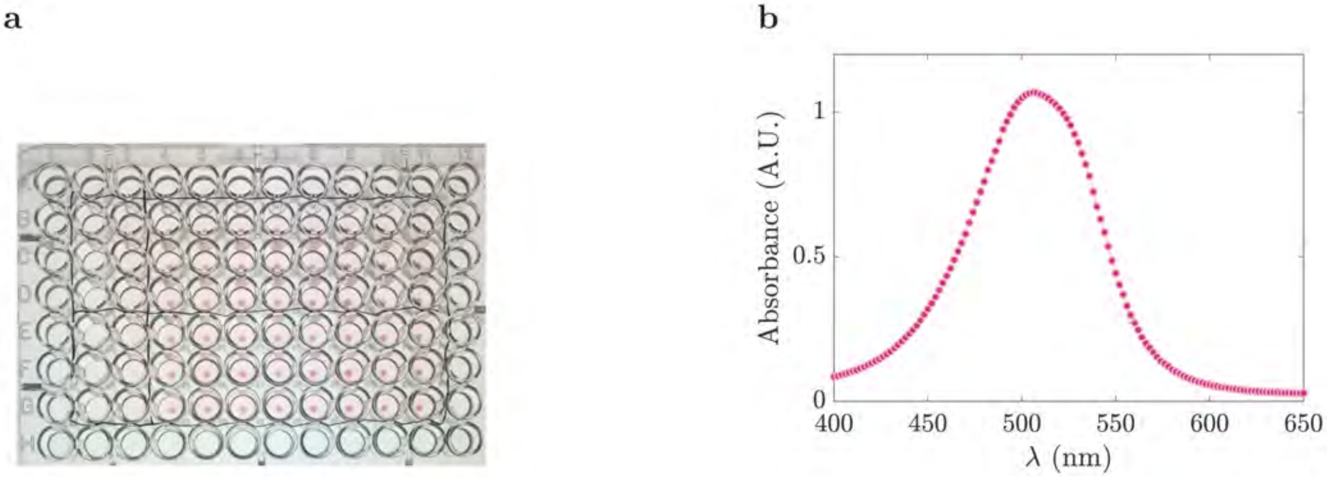


**Supplementary Figure 11. Biofilm staining with safranin O.** (a) Image of one of our 96-well plate biofilm assays, after staining with safranin O. (b) Absorption spectrum (400-800 nm, 50 nm steps) of the safranin O stain (0.1 % $v/v$ in water) diluted (1:20) in hydrochloric acid (0.1 M).
